# Simultaneous Estimation of Population Receptive Field and Hemodynamic Parameters from Single Point BOLD Responses using Metropolis-Hastings Sampling

**DOI:** 10.1101/233619

**Authors:** Stanisław Adaszewski, David Slater, Lester Melie-Garcia, Bogdan Draganski, Piotr Bogorodzki

## Abstract

We introduce a new approach to Bayesian pRF model estimation using Markov Chain Monte Carlo (MCMC) sampling for simultaneous estimation of pRF and hemodynamic parameters. To obtain high performance on commonly accessible hardware we present a novel heuristic consisting of interpolation between precomputed responses for predetermined stimuli and a large cross-section of receptive field parameters. We investigate the validity of the proposed approach with respect to MCMC convergence, tuning and biases. We compare different combinations of pRF - Compressive Spatial Summation (CSS), Dumoulin-Wandell (DW) and hemodynamic (5-parameter and 3-parameter Balloon-Windkessel) models within our framework with and without the usage of the new heuristic. We evaluate estimation consistency and log probability across models. We perform as well a comparison of one model with and without lookup table within the RStan framework using its No-U-Turn Sampler. We present accelerated computation of whole-ROI parameters for one subject. Finally, we discuss risks and limitations associated with the usage of the new heuristic as well as the means of resolving them. We found that the new algorithm is a valid sampling approach to joint pRF/hemodynamic parameter estimation and that it exhibits very high performance.

## Introduction

Modelling is an important domain of science in general and a recurring topic in population receptive field (pRF) research in particular, where functional magnetic resonance imaging (fMRI) serves as evidence acquisition method. Classical approaches such as those by (Dumoulin and Wandell, 2008) and (Kay et al., 2013) focus on point estimates of parameters in predefined models motivated by physiology and empirical evidence. In the recent work of (Zeidman et al., 2016) authors introduce the formalism of Bayesian Model Selection in order to root pRF model choices in an objective quantitative measure such as Variational Free Energy. Furthermore, the proposed formulation employs a Balloon-Windkessel model for joint estimation of pRF and hemodynamic parameters. The main limitation of this approach lies in the assumption about the form of the posterior distribution of parameters, in this case Gaussian. The method is characterized as well by high computational requirements – the reference implementation of the algorithm is reported to require about 100 seconds per voxel to converge which renders its use problematic on modern PCs with exception of high-end multi-core cluster setups (the authors give an example of 192-core cluster used to estimate 14395 voxels) or small regions of interest. A Bayesian approach using slice-sampling Monte Carlo method with fixed Hemodynamic Response Function (HRF) was recently described in (Quax et al., 2016). Similarly to the variational method the sampling approach quantifies how variable the underlying receptive field is by using the uncertainty of the posterior estimate except with the added advantage of not imposing any particular form on the posterior probability distribution. The authors underline the importance of their method’s capability to estimate variability – rendered particularly relevant by the fact that receptive fields are not rigid over time, but can change due to attention effects or task demands (Klein et al., 2014). The main contribution of our work is a new approach to Bayesian pRF model estimation combining the best characteristics of the above methods – inclusion of the Balloon-Windkessel hemodynamic model, Dumoulin-Wandell and compressive spatial summation (CSS) pRF models and using sampling for model inversion therefore not imposing any form on the posterior. Furthermore in order to address potential obstacles due to high performance requirements we introduce a novel heuristic for solving the Dumoulin-Wandell pRF model by using interpolation across a lookup table containing precomputed responses for given stimuli and a large number of predefined receptive field parameters. This enables us to massively parallelize the algorithm using a graphics processing unit (GPU) implementation of the Markov Chain Monte Carlo (MCMC) scheme. Our algorithm offers choice between existing pRF models – Dumoulin-Wandell model (Dumoulin and Wandell, 2008) and compressive spatial summation (CSS) model of pRF introduced in (Kay et al., 2013) and for BOLD generation between well-established Balloon-Windkessel model (Buxton et al., 1998; Friston et al., 2000; Irikura et al., 1994; Mayhew et al., 1998), its 3-parameter version (Stephan et al., 2007) used in the recent implementation of Dynamic Causal Modelling (DCM) in Statistical Parametric Mapping (SPM) toolbox as well as a fixed user-provided HRF. Our algorithm is presented and discussed along with introduction of QPrf – its freely available implementation in the form of a standalone toolbox (https://github.com/sadaszewski/qprf) available with source code under the terms of GNU GPLv3 license. We demonstrate CSS-pRF/Balloon-Windkessel model inversion using the new heuristic and compare it to a classical two-stage method. Furthermore, we compare different combinations of pRF (CSS, classical Dumoulin-Wandell) and hemodynamic (5-parameter and 3-parameter Balloon) models within QPrf and against existing Bayesian inversion software (BayespRF). Finally, we discuss risks and limitations associated with usage of the new heuristic as well as means of resolving them.

Visual field mapping consists of measuring responses to rings and wedges stimuli presented at varying visual field locations. Within each voxel the experimenter estimates the visual field position that produces the largest fMRI response. However, in reality the population of neurons in such voxel responds (with varying intensity) to a whole range of visual field locations. The region of visual space that stimulates the voxel is called the population receptive field (pRF) (Victor et al., 1994). The pRF method can provide estimates for receptive field location, size, orientation, laterality and surround suppression ((Kay et al., 2013; Zeidman et al., 2016)). To this end a series of stimuli is specifically designed to differentiate between the above parameters. Temporal responses are then used to fit model values with best support from the observed data (evidence).

In Dumoulin and Wandell (2008) the authors propose a quantitative approach for estimating population receptive field (pRF) parameters using a model-based coarse-to-fine optimization scheme. The pRF model is defined as two-dimensional Gaussian with means corresponding to pRF position in the visual field and a scalar covariance matrix with diagonal values equal to (pRF size)^2^. Subsequently, model parameters are varied in order to match functional magnetic resonance imaging (fMRI) time series obtained using wedges, rings and lines stimuli displayed in a series of animations. In order to do so - Frobenius inner product of stimuli and pRF Gaussian is convolved with a space-invariant hemodynamic response function (HRF) and the residual sum of squares (RSS) between the simulation and the data is iteratively minimized starting from a seed point determined by exhaustive search on predefined parameter grid.

This model is the base for further elaboration in (Kay et al., 2013) leading to the compressive spatial summation (CSS) approach. While measuring BOLD responses to a set of contrast patterns, the authors discover systematic deviation from linearity. The data are more accurately explained by a model in which a compressive static nonlinearity is applied after linear spatial summation. The authors conclude that the nonlinearity is present in early visual areas (e.g., V1, V2) and increases in anterior extrastriate areas (e.g., LO-2, VO-2). The effect of compressive spatial summation has been analyzed in terms of changes in the position and size of a viewed object. It is stated that compressive spatial summation is consistent with tolerance to changes in position and size, an important characteristic of object representation. A similar grid-based fitting approach is used for estimating parameters of the CSS-extended pRF model.

The CSS-pRF approach is characterized by simplicity and relatively good speed/accuracy of fit in most cases. Some of its shortcomings however are that it: i. provides only point estimates of the parameters; ii. does not account for spatial HRF variation (which, as acknowledged by the authors, may introduce systematic errors in pRF size estimates); iii. uses an explicit HRF model based on two gamma functions which do not allow for robust estimation of more informative hemodynamic parameters introduced by the Balloon-Windkessel model (Buxton et al., 1998).

The advancement proposed by this work is a Bayesian approach to joint estimation of pRF and hemodynamic parameters full posterior distributions using a forward signal generation model and Markov Chain Monte Carlo (MCMC) sampling. Furthermore, due to the computational costs incurred by MCMC, an optimized implementation using OpenCL is presented which allows one to take advantage of modern Graphics Processing Units (GPUs) in order to keep the processing time within the same order of magnitude as previous method while providing richer and more robust results.

## Materials and Methods

### PRF Model

A population receptive field (pRF) is the region of the visual field within which stimuli evoke responses from a local population of neurons. In (Dumoulin and Wandell, 2008) the authors proposed a model [Figure 7] of neuronal population receptive field defined by a two-dimensional Gaussian function:

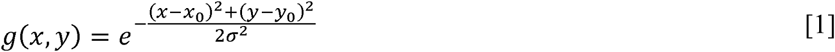

where *(x0, y0)* is the receptive field center and *σ* is the Gaussian standard deviation. Subsequently, the predicted pRF response *r(t)* is defined as sum of cells in element-wise (Hadamard) product of effective stimulus *s(x, y, t)* and the Gaussian *g(x, y)*:

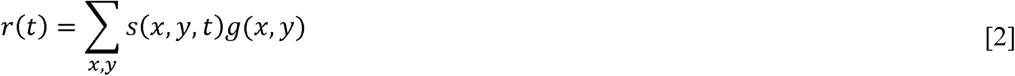

**Figure 7 (1-column).**
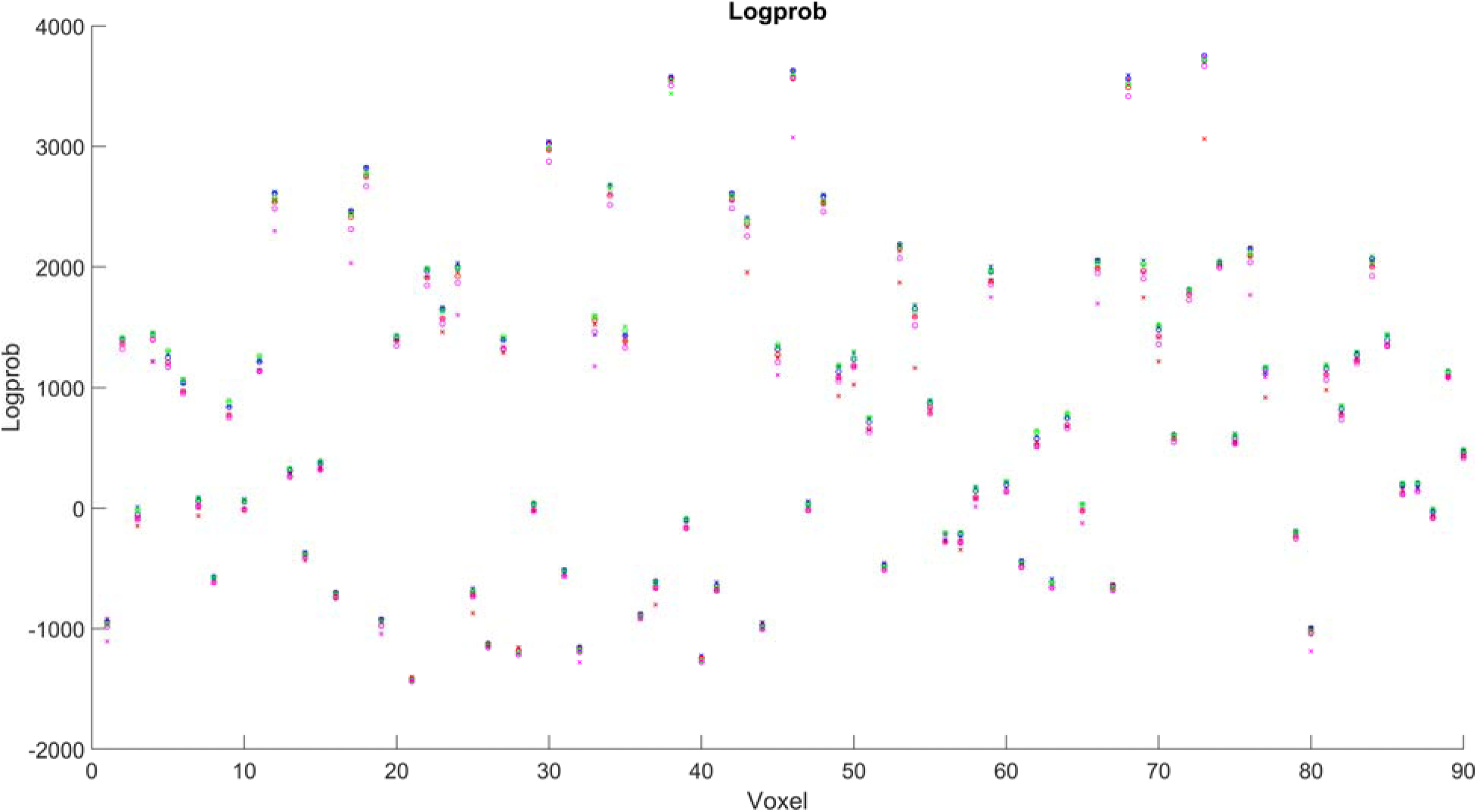
Log probability comparison between models. Blue X – noLUT_CSS_5, red X – noLUT_CSS_3, green X – noLUT_DW_5, magenta X – noLUT_DW_3, blue circle – Q_CSS_5, red circle Q_CSS_3, green circle – Q_DW_5, magenta circle – Q_DW_3.

The BOLD signal time series prediction *p(t)* is then obtained by convolving *r(t)* with a model hemodynamic response function (HRF) *h(t)*:

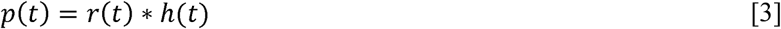

This model is further elaborated in (Kay et al., 2013) leading to compressive spatial summation (CSS) approach (Figure 1), which defines *r(t)* as:

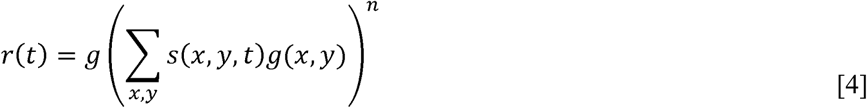

where *g* is a gain parameter and *n* is an exponent parameter. This additional compressive static nonlinearity has been proven to better explain experimental data.

**Figure 1.**
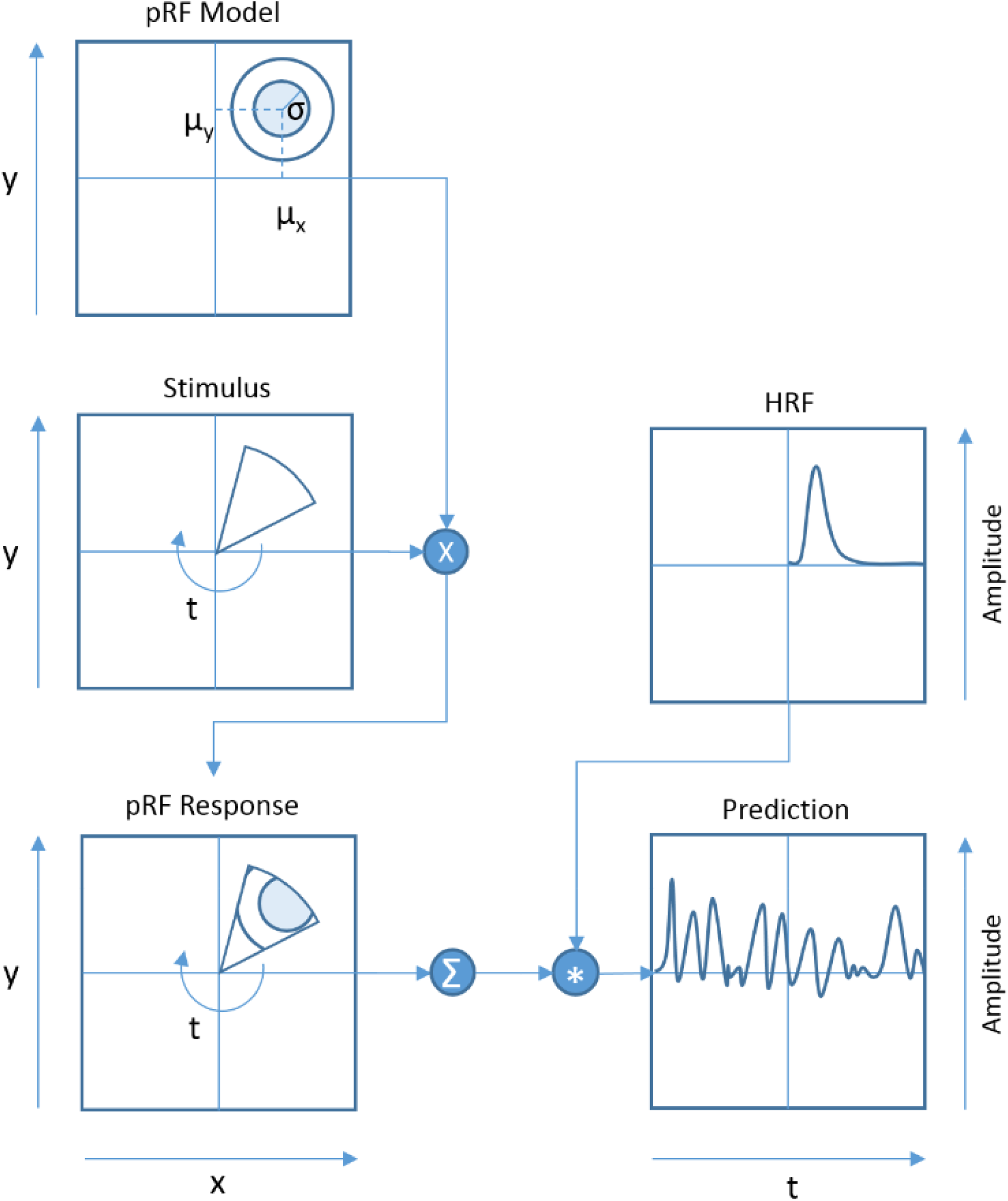
Dumoulin-Wandell pRF Model (1-column figure). Neuronal population receptive field is modelled as a two-dimensional Gaussian function (1^st^ row) where (*μ_x_*, *μ_y_*) is the receptive field center and *σ* is the Gaussian standard deviation. Two-dimensional visual stimulus (2^nd^ row, left) is multiplied element-wise with the Gaussian (3^rd^ row, left). Sum of all cells of the resulting matrix gives the pRF response. Convolution of pRF response with canonical HRF response (2^nd^ row, right) gives the final output of the model (3^rd^ row, right).

In contrast to previous studies, we use the CSS component for modeling the pRF response but instead of using convolution with a spatially invariant canonical HRF to obtain the predicted BOLD time series in [Equation 3], we employ the Balloon-Windkessel model described in the following section. We do this to account for per-voxel variability of parameters determining hemodynamic response.

### Balloon-Windkessel Model

The hemodynamic model (Figure 2) used in this study is a combination of the Balloon model and regional cerebral blood flow (rCBF) model as introduced in (Friston et al., 2000) and used for dynamic causal modelling (Friston et al., 2003). The remainder of this section contains a brief summary of the model [Figure 6].

**Figure 2.**
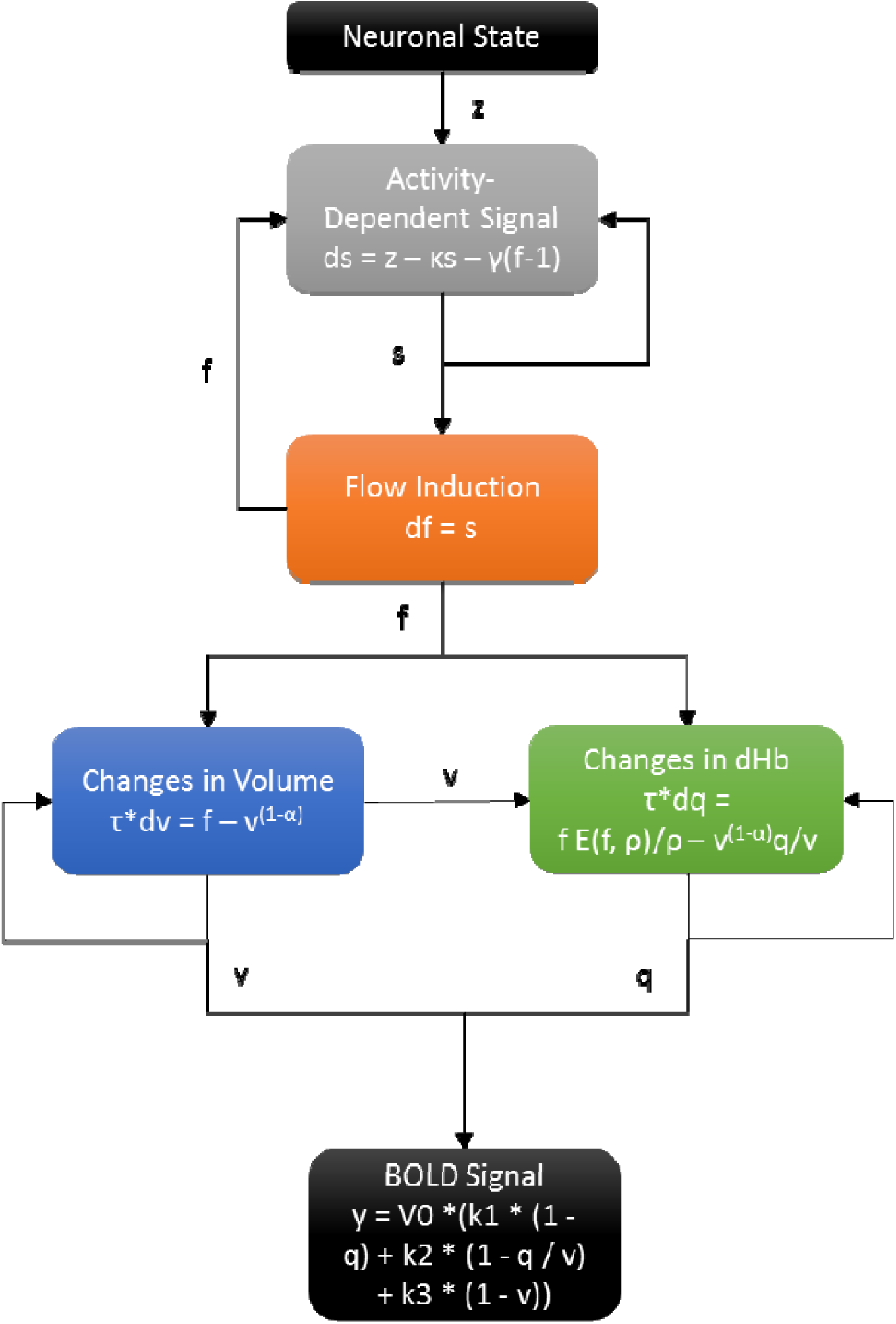
Balloon-Windkessel model (1-column figure). Relative change in BOLD signal (5^th^ row) can be expressed as a nonlinear function of normalized venous volume (*v*), normalized total deoxyhemoglobin content (*q*) and oxygen extraction fraction (*E_0_*). Changes in volume and deoxyhemoglobin content are described with coupled differential equations (4^th^ row) driven by cerebral blood flow (*f*). Flow induction is described by regional cerebral blood flow (rCBF) model (2^nd^ and 3^rd^ rows) triggered by neuronal activity (1^st^ row).

**Figure 6 (1-column).**
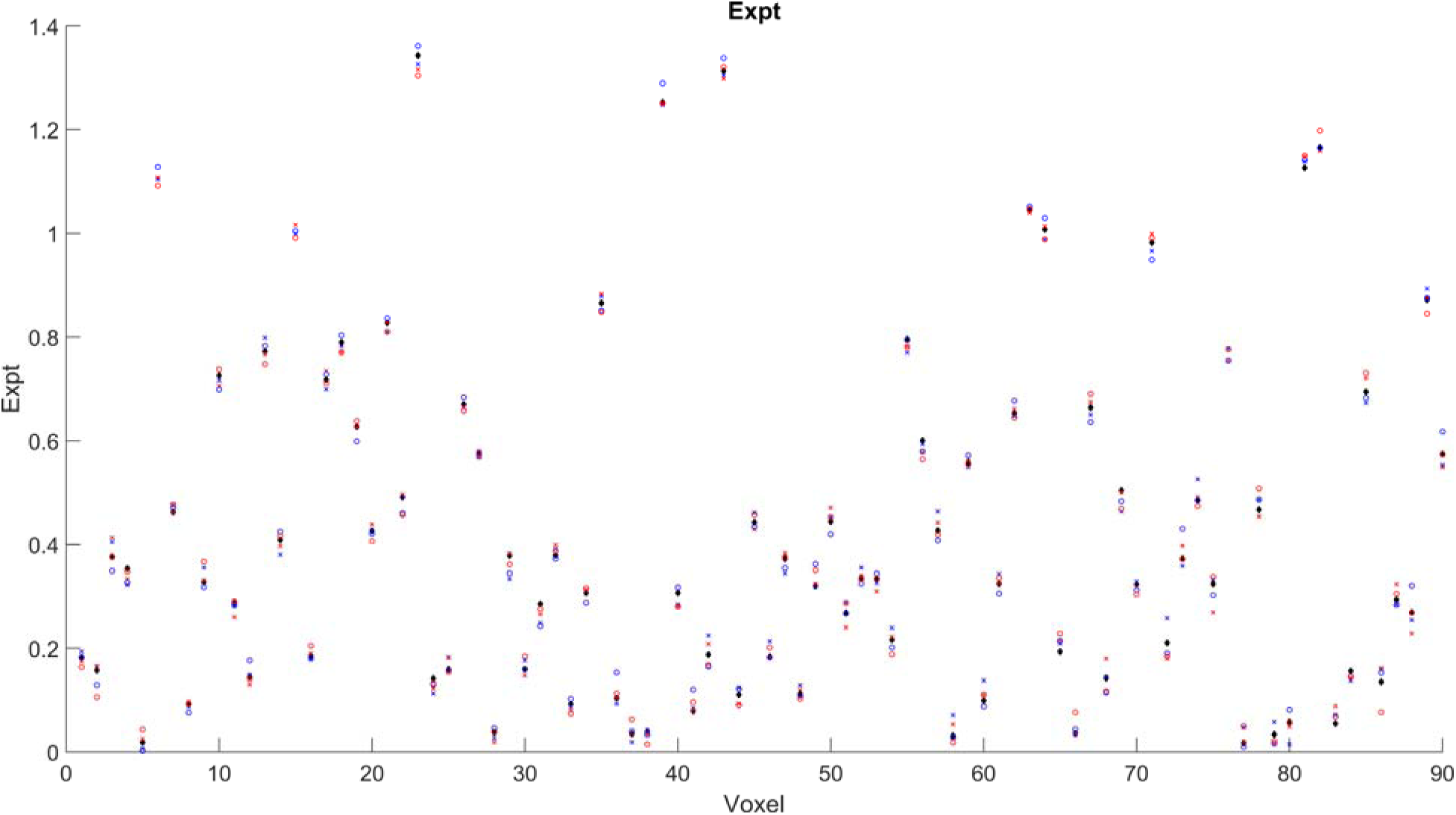
Ground truth and exponent estimates using different models for simulated data. Black diamond – ground truth. Blue X – noLUT_CSS_5, red X – noLUT_CSS_3, blue circle – Q_CSS_5, red circle Q_CSS_3.

The Balloon component is responsible for coupling rCBF to the BOLD signal (Buxton et al., 1998). The BOLD signal is modelled as the volume-weighted sum of intrinsic extravascular signal (*S_e_*) and the intravascular signal (*S_i_*):

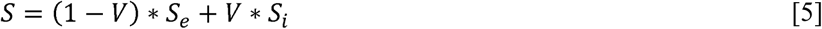

Relative change in this signal can further be expressed as a nonlinear function of normalized venous volume (*v*), normalized total deoxyhemoglobin content (*q*) and oxygen extraction fraction (*E_0_*):

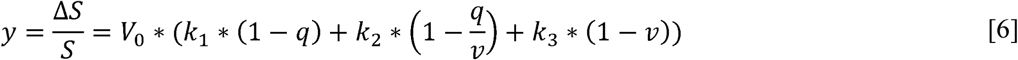

where the first term describes the intrinsic extravascular signal, the second term describes the intravascular signal, and the third term describes the effect of changing the balance of the sum in Eq. 5. *V_0_* indicates resting blood volume fraction and the *k* factors are modeled as follows:

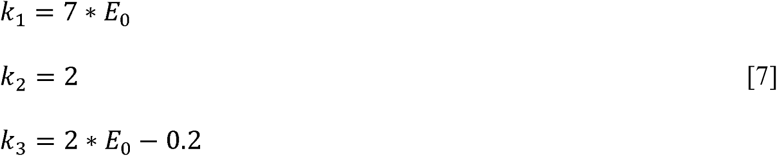

The rate of volume change *v*□ is expressed as:

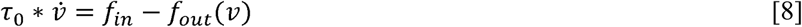

The constant *τ_0_* represents the average time it takes to traverse the venous compartment or for that compartment to be replenished (mean transit time) and is equal to *V_0_/F_0_* where *F_0_* is the resting flow. The outflow (*f_out_*) is modelled with single parameter *α*:

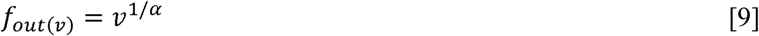

Steady-state values of *α* have been estimated by (Grubb et al., 1974) *α* ≈ 0.38 and (Mandeville et al., 1999) *α* ≈ 0.36.

Deoxyhemoglobin content change *q*□ is modelled as the difference:

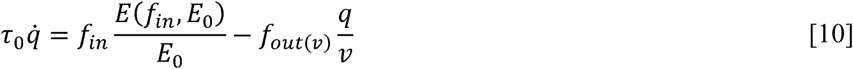

where the first term expresses the delivery of deoxyhemoglobin into the venous compartment, the second term its expulsion and *E(f_in_, E_0_)* is the fraction of oxygen extracted from the inflowing blood.

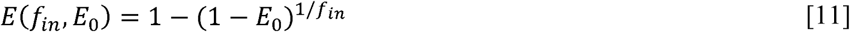

Therefore, the Balloon component contains three parameters – *E_0_, τ_0_* and *α* specifying respectively resting oxygen extraction fraction, mean transit time and stiffness exponent. Together they determine the flow-volume relationship of the venous balloon.

The rCBF component serves to specify the missing inflow state. To this end, following results from (Miller, K. L., Luh, W. M., Liu, T. T., Martinez, A., Obata, T., Wong, E. C.,…& Buxton, 2000) and (Friston et al., 2000) linear relationship between blood flow and synaptic activity is assumed resulting in a parsimonious model of inflow:

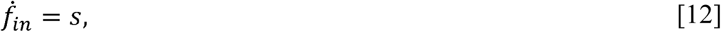

where *s* is a flow inducing signal generated by neuronal activity *u(t)* subject to:

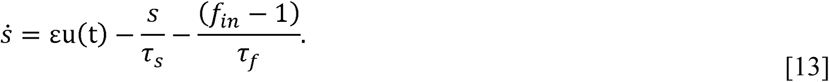

*ε*, *τ_s_* and *τ_f_* are the three unknown parameters of this component and represent respectively: the efficacy with which the neural activity causes increase in signal, the time constant for signal decay and time constant for autoregulatory feedback from blood flow (Irikura et al., 1994; Mayhew et al., 1998).

For the remainder of this work, values of *ε*=1 and *V_0_*=0.02 are assumed in line with the way this model is applied in Dynamic Causal Modelling (Friston et al., 2003) software package.

### Metropolis-Hastings Sampling

The Metropolis-Hastings (MH) scheme consists of an evolving state variable *Θ* = {*μ_x_, μ_y_, σ, κ, γ, τ, α, ρ, a, g*} and a list of all previously accepted values of *Θ.* Initial value of *Θ(t=0)* is provided by using existing point-estimate method for pRF parameters (Kay et al., 2013) and prior means for hemodynamic parameters (Empirical Bayes approach). Subsequently a proposal *Θ(t+1)* is picked in the neighborhood of *Θ(t)* using normal random distributions with small variances configurable separately for each parameter. The new value of *Θ* replaces the current one (and is added to the list) with probability *p=min(1, p(y | Θ(t+1)/p(y | Θ(t)))* which depends only on the last value of *Θ(t)* therefore satisfying the Markov criterion. As t → ∞ distribution of accepted samples approaches the desired posterior distribution *p(Θ | y)* (Hastings, 1970). The initial amount of samples *m* is skipped (this process is called burn-in period) to account for the convergence of *Θ* to the desired distribution. Furthermore, only every k-th remaining accepted sample is taken into account in the final posterior sample (this process is called thinning) to compensate for the correlation between successive random samples and ensure unbiased sampling.

The model inversion procedure relies on a stationary noise estimate obtained by a procedure loosely inspired by (Dumoulin and Wandell, 2008) and can be summarized in the following steps: 1) Principal Components Analysis (PCA) of the stimuli; 2) General Linear Model (GLM) using a subset of n=128 first principal components as independent variables and acquired Blood Oxygenation Level Dependent (BOLD) response as dependent variable; 3) 1st order autoregressive (AR) model for the residuals of the GLM model; 4) noise statistic estimation based on AR residuals; 5) response whitening by removing AR predictions; 6) a Metropolis-Hastings (MH) scheme with 3 parameters for pRF model: horizontal position (*μ_x_*), vertical position (*μ_y_*) and size (σ), 5 for Balloon-Windkessel model: rate of signal decay (*κ*), rate of flow-dependent elimination (*γ*), hemodynamic transit time (*τ*), Grubb’s exponent (*α*), resting oxygen extraction fraction (*ρ*) and 2 for compressive spatial summation which has to be applied after the Balloon-Windkessel model evaluation: exponent (a) and gain (g).

Priors (see [Table 1]) were established using an Empirical Bayes approach where applicable. The prior for the *σ* parameter was obtained by fitting a Gaussian distribution to single subject results from an existing point-estimation method (Kay et al., 2013). Non-informative priors with means at (0, 0) and large variances have been established for *μ_x_* and *μ_y_*. For hemodynamic parameters *κ, γ, τ, α, ρ* priors reported in (Friston et al., 2003) were used. Gaussian priors with large variances and means calculated from maximum likelihood fit to the whitened detrended BOLD data were used for *a* and *g* respectively.

**Table 1.** MCMC prior and proposal distributions. Note 1: the parameter space is bounded so that *μ_x_, μ_y_* and *σ* are in the range [0, 100], *a* and *g* are positive. It is achieved by rejecting samples where any of the parameters is outside of the allowed range. Note 2: Receptive field size depends on eccentricity and visual area (Amano et al., 2009; Kay et al., 2008; Larsson and Heeger, 2006; Smith et al., 2001; Winawer et al., 2010) therefore usage of an informative prior is disputable; our method readily supports using a noninformative prior instead.

| Parameter | Name | Prior (Gaussian) |  | Proposal<br>(Gaussian with mean at current point estimate) |
| --- | --- | --- | --- | --- |
|  |  | Mean | Standard Deviation | Standard Deviation |
| $\mu_x$ | Receptive field horizontal center (in pixels of stimulus mask) | 0 | Infinity | 10 |
| $\mu_y$ | Receptive field vertical center (in pixels of stimulus mask) | 0 | Infinity | 10 |
| $\sigma$ | Receptive field size (in pixels of stimulus mask) | 33.3 | Infinity | 1 |
| $a$ | CSS Exponent | 0.5 | 0.5 | 0.05 |
| $g$ | CSS Gain | 10 | 10 | 1 |
| $\kappa$ | Vasodilatory signal decay | 0.65 | 0.015 | 0.0015 |
| $\gamma$ | Rate constant for autoregulatory feedback by blood flow | 0.41 | 0.002 | 0.0002 |
| $\tau$ | Transit time | 0.98 | 0.0568 | 0.00568 |
| $\alpha$ | Grubb's stiffness exponent | 0.32 | 0.0015 | 0.00015 |
| $\rho$ | Deoxyhemoglobin content | 0.34 | 0.0024 | 0.00024 |

The summary of MCMC evaluation is contained in [Table 2].

**Table 2.**
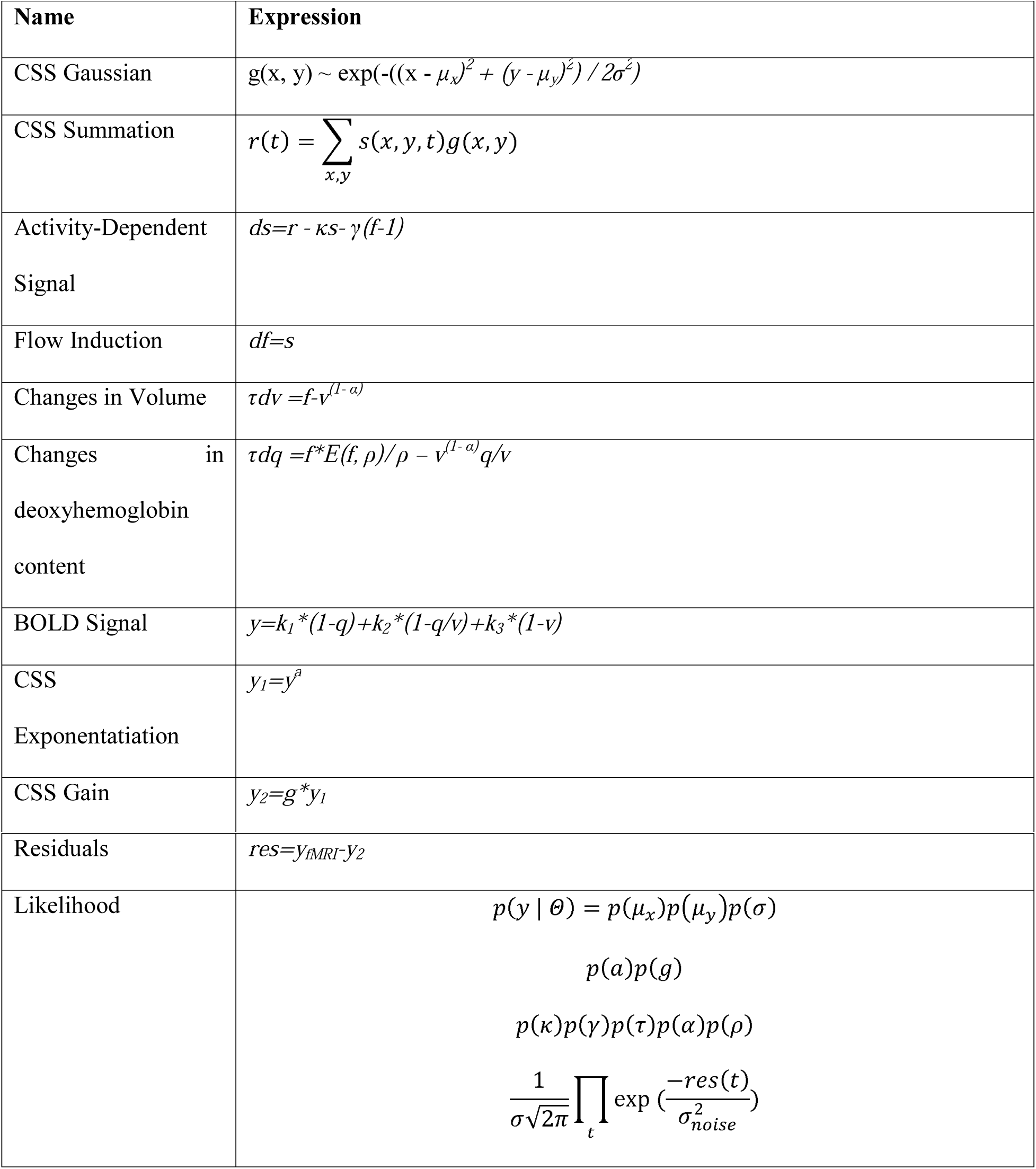
MCMC scheme summary

### Algorithm Listing Conventions

The following two sections describe the main CPU and GPU algorithms employed by our implementation of the estimation procedure. We assume the following conventions in the listings below. Variables are written in *italics*, constants are CAPITALIZED, pseudo-code keywords are written in **bold**.

Two distinct notations are used for describing ranges of values. The bracket notation [start, end) signifies a set of values {*start, start + 1,*…, *end − 2, end − 1*}. The colon notation [*start:step:end*) signifies the following set of values: {*start, start + step, start + 2 * step,*…, *start + (k - 1) * step, start + k * step*} where *k* is the biggest integer such that *start + k * step < end*. Sets are treated as row vectors where relevant.

The pseudecode refers to a number of built-in functions. bsxfun(*fun, A, B*) is inspired by a MATLAB routine with the same name. It takes a lambda expression *fun* and applies it to corresponding pairs of elements from *A* and *B*. Implicit expansion is used in case the shapes of *A* and *B* don’t match exactly. The term “implicit expansion” signifies that the smaller array is repeated across singleton dimensions in order to match the shape of the bigger one.

Routine pinv(*A*) computes a Moore-Penrose pseudoinverse of matrix *A*. The computation is based on singular value decomposition (SVD) algorithm with automatically adjusted tolerance *tol* = max(*ncols, nrows*) * abs(*sv*[0]) where *ncols* and *nrows* signify respectively number of columns and rows in the matrix and *sv* is an array of singular values in descending order of absolute magnitude.

Function *repmat(A, {m, n})* repeats matrix *A m* times vertically and *n* times horizontally.

Furthermore, for reason of brevity some of the convenient MATLAB syntax is employed in listing of Algorithm 2. Expression *“@power”* signifies lambda function raising its first argument to the power expressed by the second argument. Expression *A*’ signifies transpose of a matrix.

Standard trigonometric functions *sin(a)* and *cos(a)* compute respectively sine and cosine of an angle *a* expressed in radians. The function *min(a, b)* returns the smaller of two scalar values *a* and *b*.

Notation *N(A)* is a highly abbreviated notation used to signify calculation of element-wise log-probabilities using Gaussian distribution with mean and standard deviation corresponding to prior distribution characteristic for quantity in array *A*.

Notation *randn(…)* is used to signify generating random values according to normal distribution with zero mean and standard deviance corresponding to step size selected for respective components of MH state variable.

Function *uniform_random(a, b)* returns a random real number from range [*a*, *b*].

### CPU Algorithms

The Metropolis-Hastings sampling (Algorithm 1) constitutes core of the implementation. It is divided into Initialization, Data Preprocessing and Sampling phases. The Sampling phase is further divided into GPU part and CPU part depending on the device performing bulk of the computation involved and Stepping part responsible for performing the random walk.

Initialization phase loads acceleration lookup table (*stage2*) and initial pRF parameter estimates (*xs, ys, rfsizes, expts, gains*), prepares polynomial detrending matrix (*pmatrix*) and bootstraps remaining working variables.

In Data Preprocessing phase, MRI data are divided into batches and processed using preprocess_data() GPU routine (Kernel 5).

Subsequent Sampling phase runs a predefined number of iterations (*NumberOfSteps*) of a loop consisting of GPU and CPU computation. The GPU part is responsible for executing the signal generation model in three steps: reset() routine (Kernel 2) restores initial states of the Balloon-Windkessel model variables, fwd_model() kernel (Kernel 3) performs a forward run of combined pRF and Balloon-Windkessel MRI signal generation model, whereas eval_proposal() function (Kernel 6) computes the logprobability of resulting signal given registered MRI data (evidence). The CPU part adds logprobabilities resulting from priors and for each voxel decides whether to accept state variable values or revert to old ones. The latter is done in stochastic manner by taking ratio of current state probability to previous state probability and comparing against a realization of uniformly random variable from range [0, 1]. If the random value is less, new sample is accepted, otherwise it is rejected. If new state has higher probability than the old state, it will always be accepted because ratio is larger than one. Accepted samples are added to respective per-voxel sets of samples for individual state variables. Finally, the next proposal state is generated by adding normal random variable with zero mean and predefined standard deviation to current values of each state variable.

#### Algorithm 1.

*Metropolis-Hastings sampling* – Perform posterior sampling on joint pRF / Balloon-Windkessel model

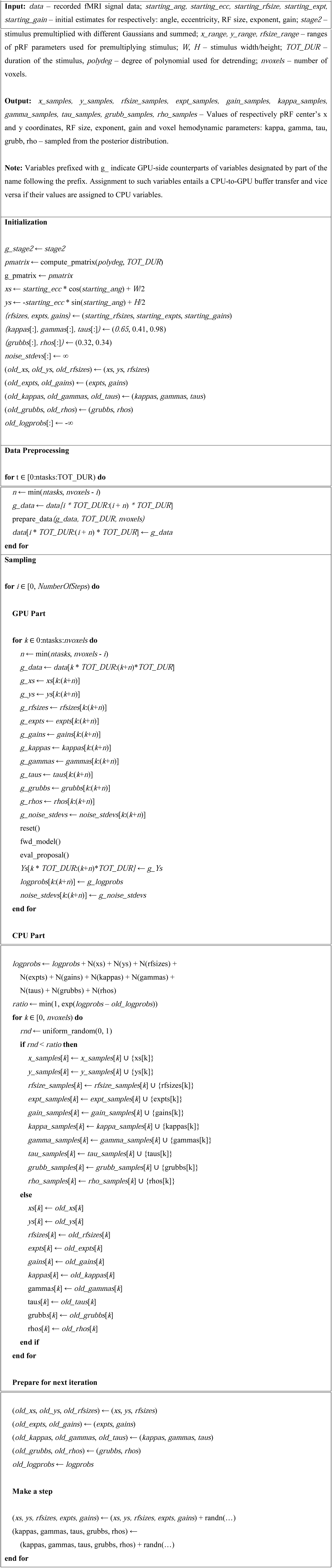

The compute_pmatrix() routine (Algorithm 2) computes polynomial detrending matrix. To this end, a matrix of the form:

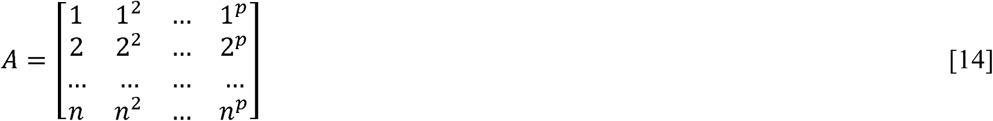

is created where n is the duration of the MRI stimuli/signal and p is the chosen polynomial degree. Then, the expression:

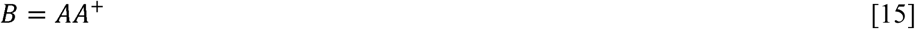

is the polynomial detrending matrix where A^+^ signifies Moore-Penrose pseudoinverse of matrix A.

#### Algorithm 2.

*compute_pmatrix()* – Compute polynomial detrending matrix

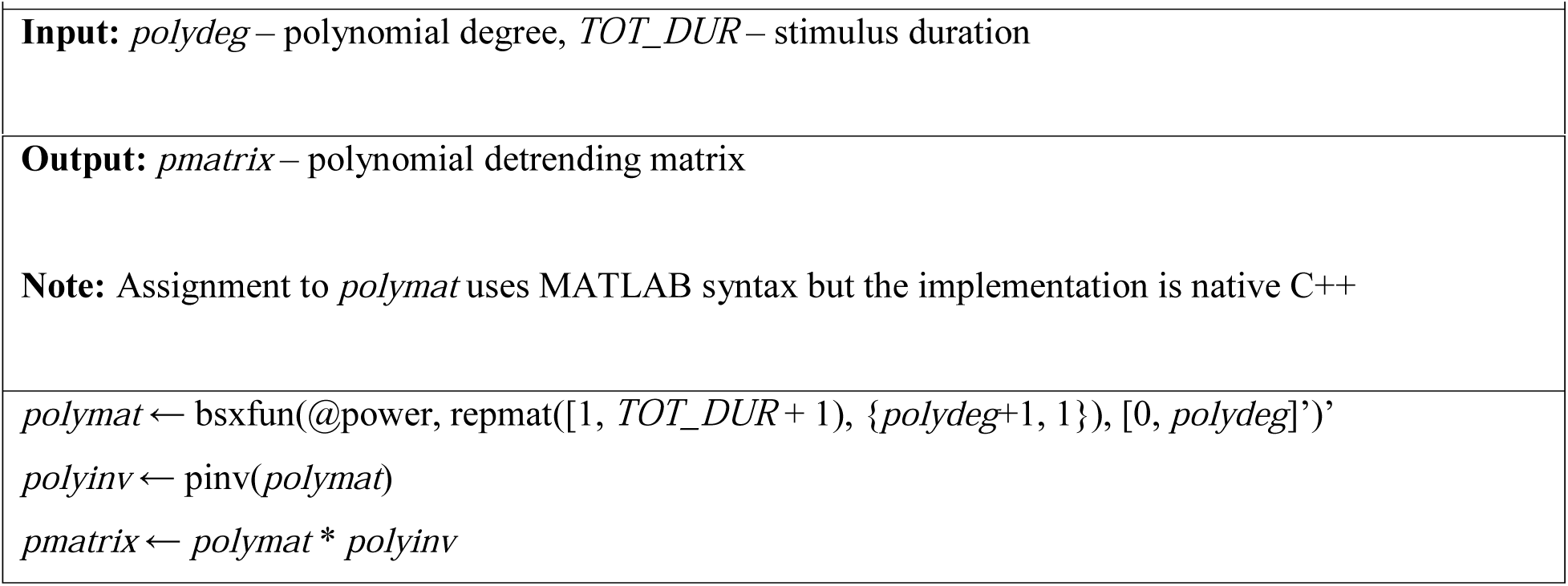

### GPU Algorithms

The key observation that allowed acceleration of the sampling procedure was that the 2D visual stimuli is uniform across all voxels and can be premultiplied and summed over a wide range of Gaussians to create a lookup structure (Figure 3) which can then be used to approximate results of such operation for *any* Gaussian we encounter during the random walk by taking eight closest precomputed signals and performing a trilinear interpolation. This function is performed by the interp_gaussian() GPU routine (Kernel 1).

#### Kernel 1.

*interp_gaussian()* - Interpolate premultiplied stimuli according to specified pRF parameters

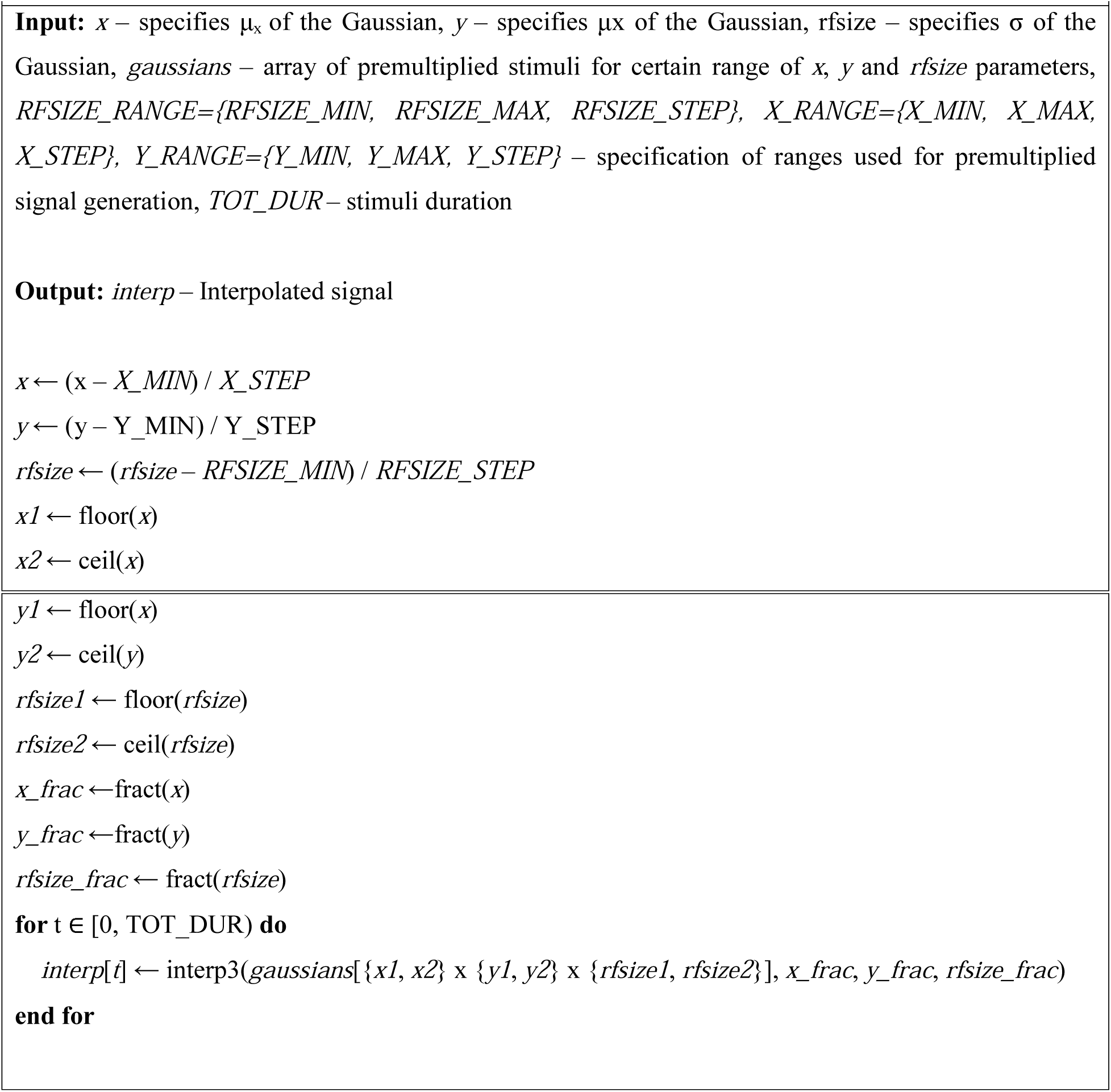

**Figure 3.**
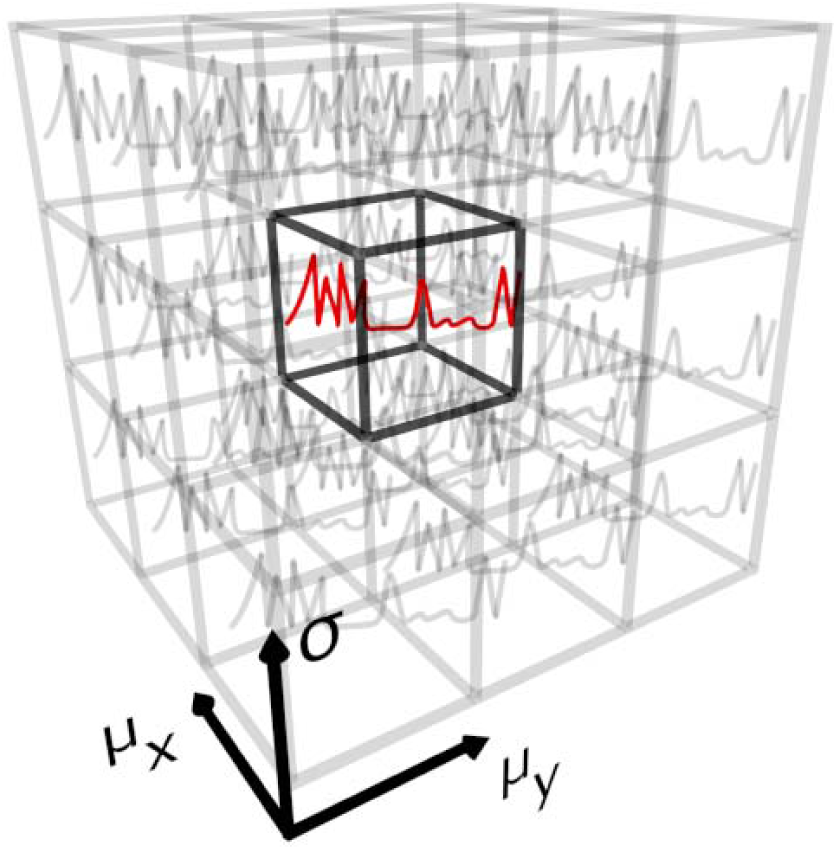
Premultiplied signal illustration (1-column figure). 2D visual stimuli is the same for all voxels so it can be premultiplied and summed over a wide range of Gaussians to create a lookup structure valid across the entire dataset. For arbitrary values of μ_x_, μ_y_, σ eight surrounding cells from the lookup table are picked and the resulting signal is approximated using trilinear interpolation. This scheme implicitly bounds the pRF model parameter space as the Gaussian parameters cannot exceed the bounds of the lookup structure.

The joint pRF / Balloon-Windkessel model is handled by the following three GPU procedures. The reset() call (Kernel 2) restores initial values of Balloon-Windkessel state variables.

#### Kernel 2.

*reset()* – Reset simulation data to baseline values

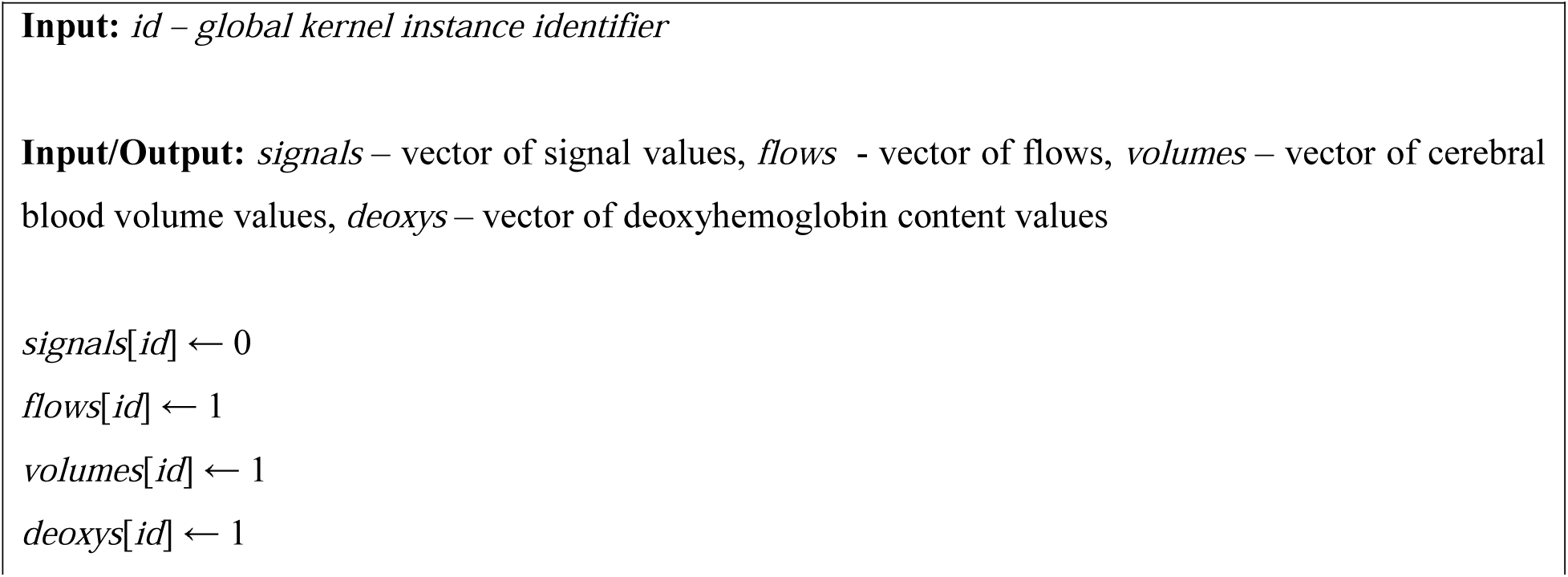

The fwd_model() function (Kernel 3) initializes variables pertaining to the joint model and uses interp_gaussian() to obtain stimuli multipled with Gaussian with parameters for current random walk state. Such signal is used as the neural activity input for the Balloon-Windkessel model. Subsequently evaluation of the latter is performed over the time corresponding to one stimulus frame. The output of Balloon-Windkessel model is then subjected to remaining part of the compressive spatial summation model, namely exponentiation and multiplication by the gain factor to account for unknown MRI signal units. Results are detrended using polynomial detrending matrix (*pmatrix*). Finally, Balloon-Windkessel model state is updated for current voxel and output copied from local array to global buffer accessible from CPU.

#### Kernel 3.

*fwd_model()* – Perform joint pRF / Balloon-Windkessel model simulation

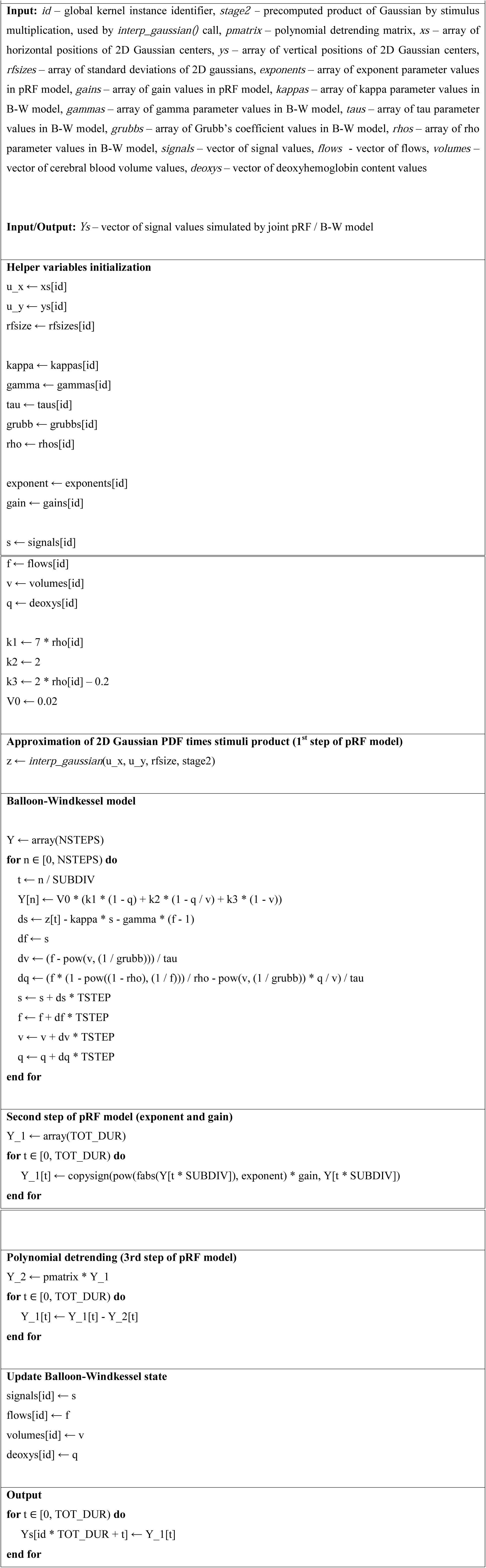

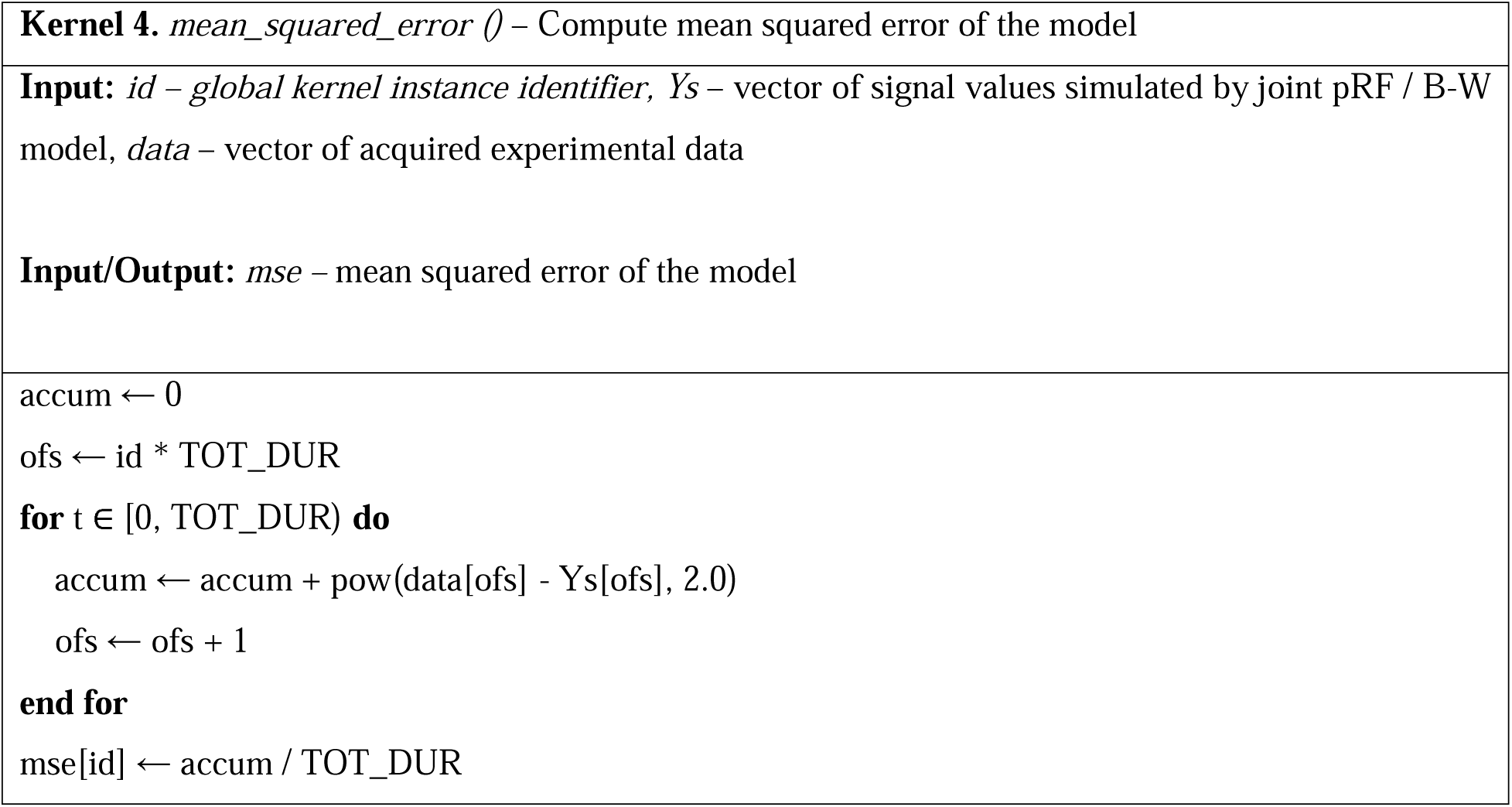

#### Kernel 5.

*prepare_data ()* – Perform polynomial detrending on evidence data

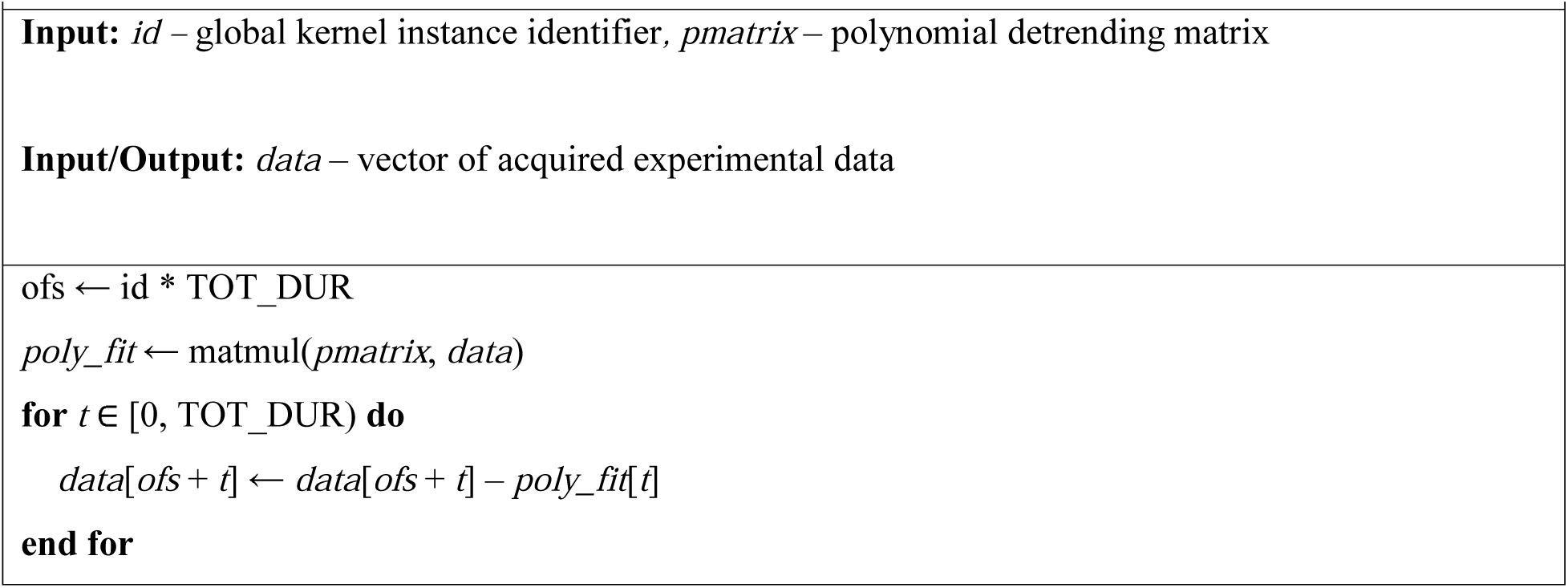

#### Kernel 6.

*eval_proposal ()* – Compute logprobability of simulated signal given evidence

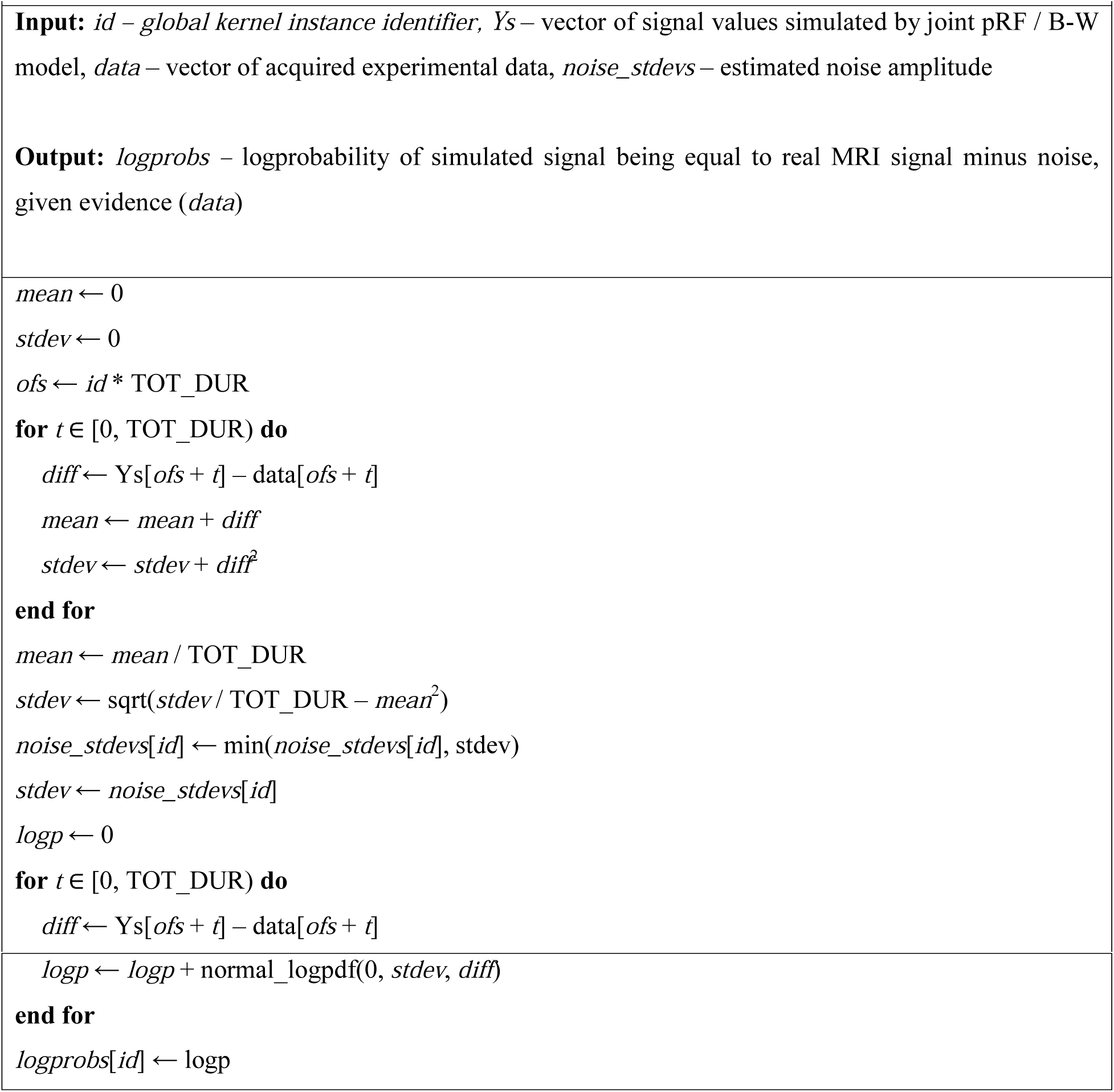

### Stimulus

Visual stimuli were created using extensions from the PsychToolbox (Brainard, 1997; Pelli, 1997) within the Matlab programming environment. An LCD projector was used to image the stimuli onto a back projection screen within the bore of the magnet. Subjects viewed the display through an angled mirror. The maximal stimulus radius was 9.15° of visual angle. Each stimulus image was at a resolution of 768 x 768 pixels. Binary stimuli masks were defined at time points which coincided with the slice-time corrected fMRI data and down sampled to a resolution of 100x100 pixels to improve efficiency of the MCMC sampling routines. For all experiments a small dot at the centre of the image served as a fixation point (4 pixel diameter) which changed color at randomized intervals between 1 and 5 seconds. Subjects were asked to fixate on the dot and indicate each time the dot changed color using an MRI compatible button box. The stimuli consisted of achromatic contrast patterns (spatially pink noise) overlaid with randomly positioned and scaled visual object stimuli from Kriegeskorte et al. (2008). The full stimuli patterns can be found at http://cvnlab.net/analyzePRF/.

In the first set of experiments the stimulus consisted of a combination of rotating wedge, contracting ring and sweeping bar stimuli. The stimuli can be broken down into the following temporal structure:

- Wedge stimuli covering a 90° angle swept 8 full counter-clockwise rotations. The wedge took 32 seconds to complete one full 360° rotation.
- Bars with a width equal to 12% of the stimulus diameter were swept across 8 unique (cardinal and oblique) directions. Each bar sweep took 32-seconds to complete (28-second sweep followed by a 4-second rest).
- Contracting ring stimuli were swept across the visual field over the course of 32-seconds (28-second sweep followed by a 4-second rest). This was repeated 5 times. The widths of the rings were scaled linearly with eccentricity to compensate for cortical magnification in visual cortex (Cowey and Rolls, 1974; Rovamo and Virsu, 1979).
- 16 second rest periods with mean luminance (zero contrast) images were placed at the start, between stimuli types and at the end of the stimulus (taking into account the additional 4 second rest periods at the end of bar and ring sweeps).

The stimulus took a total of 12 minutes 20 seconds to complete (16 + 8*32 + 16 + 4*32 + 12 + 4*32 + 12 + 5*32 + 12 s).

### In-Vivo Datasets

All MRI data was acquired with a 3T Siemens Prisma MR system using a 64-channel head coil. One male subject with no history of eye disease (aged 28) underwent all of the fMRI acquisition protocols. Foam padding and audio headphones minimized head motion. The functional MRI data were acquired using a 2D-EPI sequence (TR/TE = 1518/30 ms, 23 slices, 3x3x2.5 mm 3 resolution, 20% distance factor, FOV = 192x192 mm 2, 12% phase oversampling). The slices were oriented parallel to the calcarine sulcus, covering the majority of occipital cortex. A whole brain 2D-EPI sequence with 49 slices and 5 volumes was acquired to aid with structural registration. All other parameters of the whole brain sequence were the same as in the fMRI acquisition. A T1-weighted anatomical image (TR/TE = 2000/2.93 ms, 1x1x1 mm 3 resolution) was acquired and used to reconstruct the cortical surfaces. In order to estimate local field distortions we acquired a B0 field map (TR/TE = 1020/10 ms, 3x3x3 mm 3 resolution). Functional scans for our dataset used the combined rotating wedge, contracting ring and sweeping bar stimuli and were acquired over 488 volumes. A total of 3 fMRI scans were performed in the session and later used to investigate scan-rescan reproducibility of the estimated pRF parameters.

### Image Preprocessing

The first 5 volumes of each fMRI run were discarded to allow magnetization to reach a steady state. Preprocessing of the fMRI data was performed within SPM12 using the following procedure. Slicetime correction was applied to adjust for differences in slice acquisition times. The fMRI data was aligned to correct for movement artifacts and co-registered with the whole brain reference data. The fMRI and reference data were spatially unwarped (Andersson et al., 2001; Jezzard and Balaban, 1995) to improve coregistration between the EPI and structural data (Hutton et al., 2002). The reference data was then coregistered with the T1-weighted structural image and the same rigid-body transformation applied to the fMRI data. Binary stimuli masks were defined at time points which coincided with the slice-time corrected fMRI data and down sampled to a resolution of 100x100 pixels to improve efficiency of the nonlinear optimization routines.

The T1-weighted anatomical images were processed with FreeSurfer (Dale et al., 1999) for white and grey matter segmentation and cortical surface reconstruction. FreeSurfer’s Desikan-Killiany atlas (Desikan et al., 2006) was used to select cortical regions within the occipital lobe and to reduce to the amount of cortex over which we fit the pRF models. The preprocessed fMRI data were sampled at distance corresponding to 20% of cortical thickness along normals of FreeSurfer’s fsaverage surface. All CSS-pRF model estimations were performed on the surface projected fMRI responses. Visual field delineation was performed manually using the polar angle maps projected onto inflated cortical surfaces (Wandell et al., 2007).

### Experiments

To fully evaluate the impact of introducing the lookup table heuristic we have performed an exhaustive range of tests on simulated data and concluded with a demonstration of the new algorithm as applied to a whole-ROI in-vivo dataset.

We performed a test of the lookup table signal reconstruction accuracy compared to full pRF model evaluation. To this end we computed the root mean squared error (RMSE) of the sum of the product of the stimulus and the Gaussians with different positions and sizes. We generated 1000 Gaussians with randomized parameters and computed two signals – i. by using the full pRF model evaluation (the reference signal) and ii. by using the lookup table. We compared the RMSE between the two relative to the reference signal amplitude.

We performed a series of MCMC runs on the simulated data in order to reliably compare estimated means with the known ground truth. To this end, we generated 90 time series with pRF parameters spread uniformly across the available space and hemodynamic parameters picked randomly according to their empirical prior distributions. We added noise to the resulting signals using contrast-to-noise ratio (CNR) typical for fMRI studies – 5:1. We used randomized starting points in the MCMC scheme and used a time limit of 3 hours of sampling. We didn’t apply any burn-in or thinning, instead we report mESS measures along with the parameter estimates.

In order to add weight to our findings we have executed the above MCMC scheme for different combinations of hemodynamic (5-parameter Balloon and 3-parameter Balloon) and pRF models (CSS and classical Dumoulin-Wandell). Furthermore, we have performed the above using implementations with and without the usage of the lookup table. Therefore in total we have evaluated 8 variants of the above MCMC scheme – QPrf with LUT using CSS-pRF and 5-parameter Balloon model HRF (Q_CSS_5 and Q_CSS_5_bis, two runs to assess convergence), QPrf with LUT using Dumoulin-Wandell pRF and 5-parameter Balloon model HRF (Q_DW_5), QPrf with LUT using CSS-pRF and 3-parameter Balloon model HRF (Q_CSS_3), QPrf with LUT using Dumoulin-Wandell pRF and 3-parameter Balloon model HRF (Q_DW_3), QPrf without LUT using CSS-pRF and 5-parameter Balloon model HRF (noLUT_CSS_5 and Q_CSS_5_bis, two runs to assess convergence), QPrf without LUT using Dumoulin-Wandell pRF and 5-parameter Balloon model HRF (noLUT_DW_5), QPrf without LUT using CSS-pRF and 3-parameter Balloon model HRF (noLUT_CSS_3), QPrf without LUT using Dumoulin-Wandell pRF and 3-parameter Balloon model HRF (noLUT_DW_3).

In order to put the LUT approach to test within another sampling framework we decided to use RStan (Carpenter et al., 2017). Due to excessive run time we performed a test on a single voxel of simulated data using RStan’s NUTS sampler without using the LUT and using the LUT. NUTS settings included 1000 leapfrog iterations of warmup and 2000 leapfrog iterations of sampling. We used the same Bayesian model and the same priors as in QPrf and compared the estimation results to one another and the ground truth. We report as well differences in run time.

Finally, we applied accelerated QPrf model with CSS-pRF component and 5-parameter Balloon model to estimate all 50000 voxels in the visual ROI of the brain of one subject. We present parameter maps for pRF parameters and report our findings with respect to covariance between parameters.

## Results and Discussion

### Simulated Data Results

Figure 4 illustrates the relative RMSE error between the signal obtained from the lookup table and the signal generated using full evaluation of the model for 1000 Gaussians with randomly picked parameters. Mean RMSE relative to amplitude was equal to 2%. There were 22 points (2%) exceeding relative error of 3% with the maximum error below 5%.

**Figure 4.**
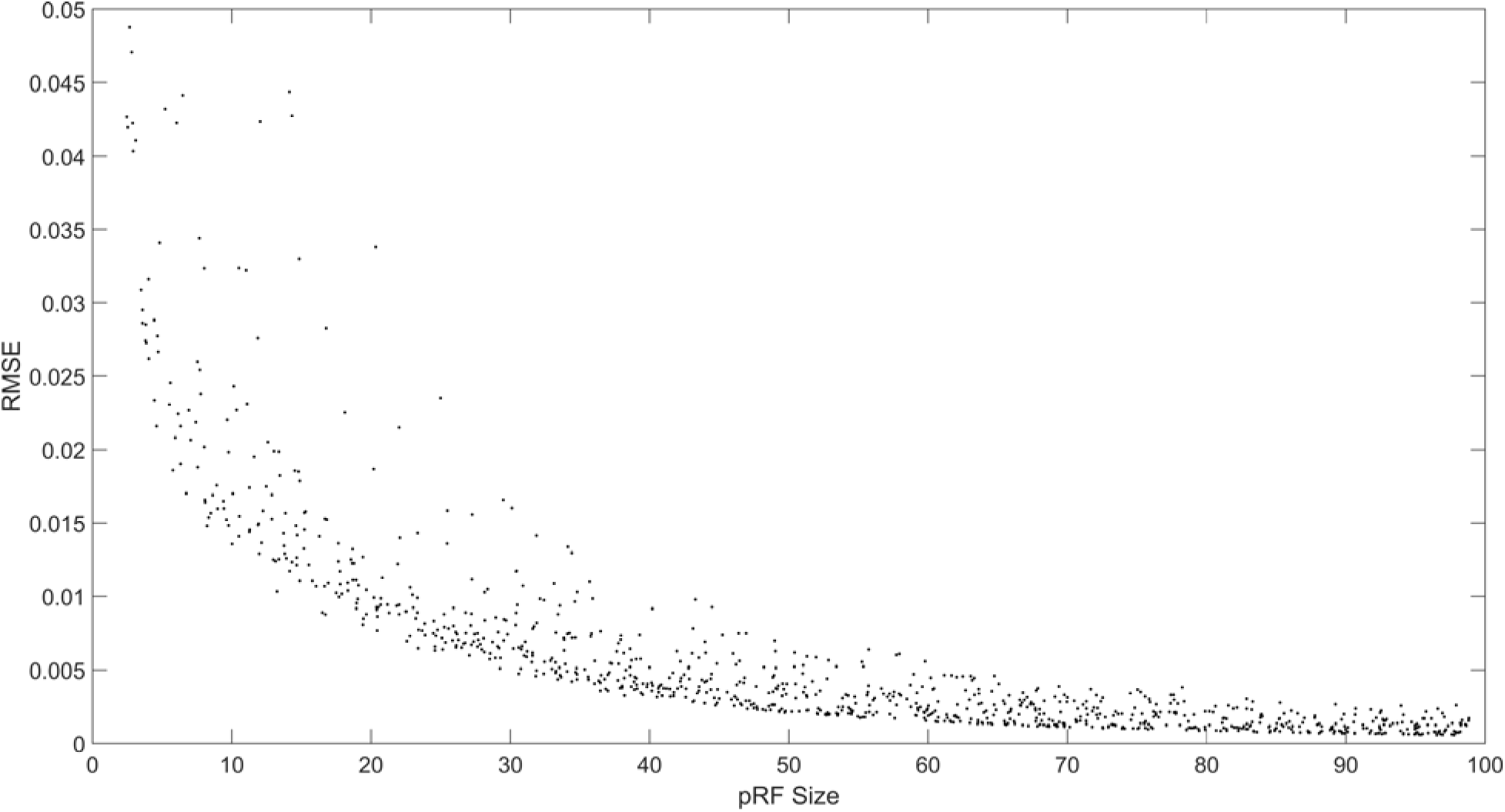
Root mean squared error (RMSE) of the sum of the product of the stimulus and the Gaussians with different positions and sizes relative to maximum signal amplitude. The figure illustrates the relative RMSE error between the signal obtained from the lookup table and the signal generated using full evaluation of the model for 1000 Gaussians with randomly picked parameters. Mean RMSE relative to amplitude was equal to 2%. There were 22 points (2%) exceeding relative error of 3% with the maximum error below 5%.

Across the parameter estimates for simulated data we observed differing degree of correlation between RF Size, Exponent and Gain. The max(p-value) (Friston et al., 1999) approach revealed significant correlation between gain and exponent. While this is not in line with the “ground truth” of the generation method were both values were picked independently - it is not a surprising result. Both exponent and gain affect the scaling of the signal. In the original compressive pRF model which included bringing simulated signal to unit length they were intended to influence the “speed” of signal edges. In our simulation and MCMC method we rely on model of physical fMRI signal in which the influence of gain and exponent seems to be largely interchangeable. The role of RF-Size might be the following: as the standard deviation of correctly centered Gaussian increases, a growing portion of the probability mass is placed outside of the considered stimulus area (100x100 in scaled stimulus space). This results in the need to increase estimates of gain and/or exponent to maintain the observed signal amplitude. This highlights the intrinsic imperfections of the model. We found no other correlations in the estimates.

### Model Comparison Results

We report the most strict (choice of batch size giving the smallest result) multivariate ESS (Vats et al., 2015) measures for all runs (Table 3). In a single plot (Figure 5) we present estimated mean comparisons between noLUT_CSS_5, noLUT_CSS_3, noLUT_DW_5, noLUT_DW_3, Q_CSS_5, Q_CSS_3, Q_DW_5 and Q_DW_3 and the ground truth. In (Figure 6) we show estimated means and ground truth for the exponent which is present only in noLUT_CSS_5, noLUT_CSS_3, Q_CSS_5 and Q_CSS_3 models. Furthermore we report log probability estimates for all methods (Table 4) and present a per-voxel plot (Figure 7).

**Table 3.** Multivariate Effective Sample Size (mESS) measures for runs of different pRF/HRF models

| HRF Model | pRF Model | With Lookup Table |  | Without Lookup Table |  |
| --- | --- | --- | --- | --- | --- |
|  |  | Mean | Stdev | Mean | Stdev |
| **3-param Balloon** | CSS | 2703 | 556 | 2239 | 355 |
| **3-param Balloon** | DW | 2196 | 391 | 2570 | 418 |
| **5-param** | CSS | 1847 | 482 | 2756 | 354 |
| **Balloon** |  |  |  |  |  |
| **5-param Balloon** | DW | 2265 | 210 | 2942 | 216 |

**Figure 5 (2-column).**
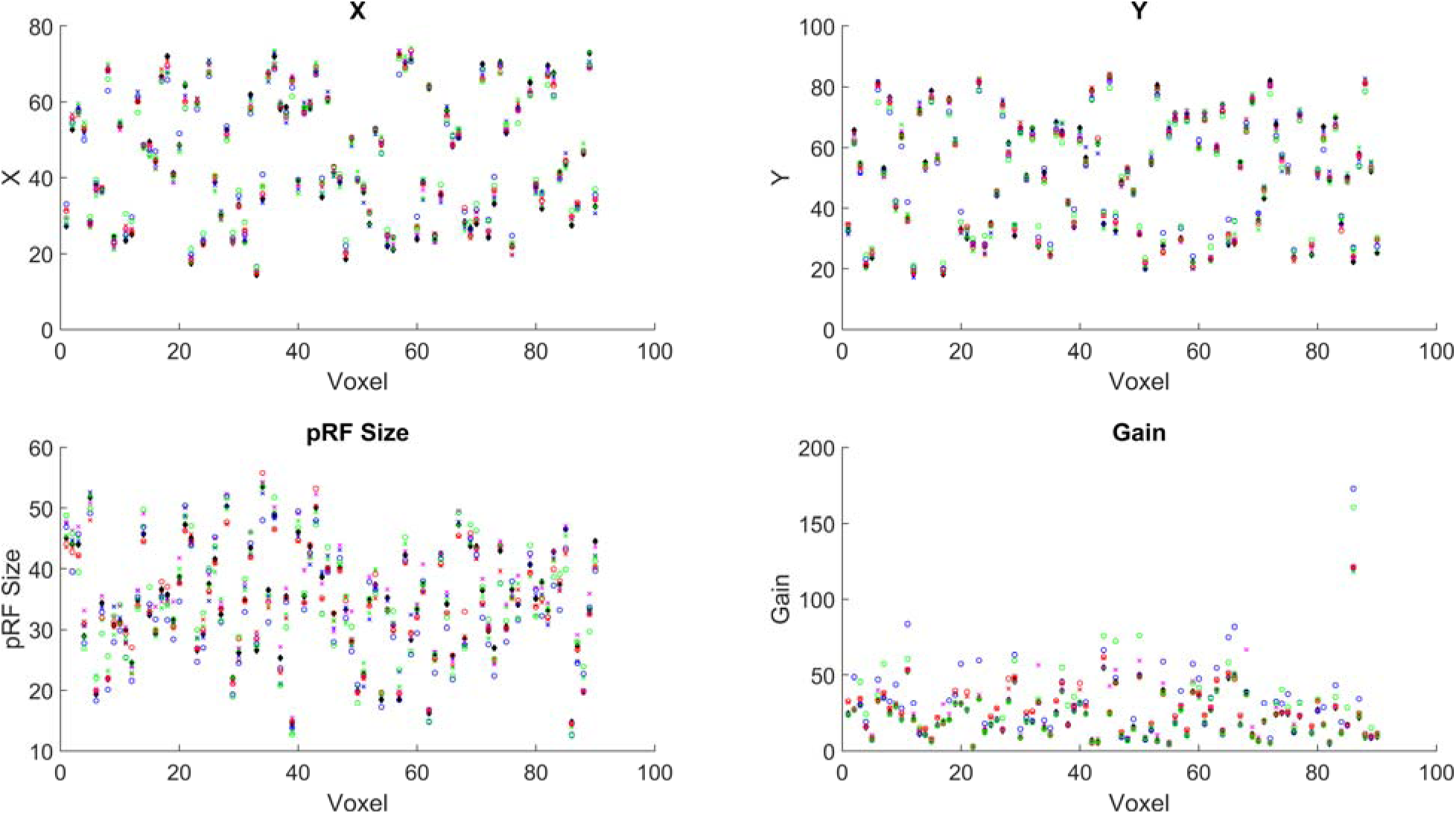
Ground truth and parameter estimates using different models for simulated data. Black diamond – ground truth. Blue X – noLUT_CSS_5, red X – noLUT_CSS_3, green X – noLUT_DW_5, magenta X – noLUT_DW_3, blue circle – Q_CSS_5, red circle Q_CSS_3, green circle – Q_DW_5, magenta circle – Q_DW_3.

**Table 4.** Log probability comparison between different pRF/HRF models within QPrf with and without the usage of the lookup table.

| HRF Model | pRF Model | With Lookup Table | Without Lookup Table |
| --- | --- | --- | --- |
| **3-param Balloon** | CSS | 665 | 360 |
| **3-param Balloon** | DW | 370 | 242 |
| **5-param Balloon** | CSS | 1122 | 1207 |
| **5-param Balloon** | DW | 1140 | 1152 |

The mESS measures obtained (Table 3) were satisfactorily large. Convergence between two runs of Q_CSS_5 was almost perfect. PRF parameter estimates common to all models (Figure 5) were largely consistent across models. We observed bigger discrepancies in pRF size and gain estimates. Position estimates were identical up to a few pixels (in rescaled – 100x100 stimulus coordinate system). Exponent estimates were also largely consistent (Figure 6) with no evidence of systematic over-/underestimation in any particular model. Hemodynamic estimates were not deviating significantly from their prior means and no systematic bias was observed. Log probabilities (Table 4, Figure 7) were higher for 5-parameter Balloon model both with and without the lookup table which is unsurprising since this was the model used to generate the data. Log probability was of similar magnitude with and without the lookup table.

### fMRI Results

We have estimated means, variances and confidence intervals for each parameter i. directly from parameter samples and ii. by fitting multivariate Gaussian distribution as the posterior density model p(Θ | y) for whole-state samples. Fitted Gaussian means were within 1SD (computed directly from samples) of the sample-based mean for 90% of the voxels, supporting Gaussian character of the distributions.

Furthermore we computed pairwise correlation coefficients using direct samples approach for each pair of model parameters. Patterns of correlation covering the whole [-1, 1] range have been observed with most points presenting close to zero correlation for all pairs of parameters. Spatial distribution of correlation coefficients didn’t exhibit any meaningful pattern suggesting that higher absolute values of correlation coefficient should be attributed to estimation uncertainty (within our model the shape of time series matching observed time series might be achieved by varying either pRF or hemodynamic parameters) rather than any systematic change of pRF / hemodynamic configuration depending on region. This stays in line with (Dumoulin and Wandell, 2008) who quote hemodynamic parameters as sources of additional uncertainty in pRF parameter estimation.

In (Figure 8) we present a whole-ROI map of pRF parameters for a single subject. The spatial distribution of angle and eccentricity is in line with literature (Dumoulin and Wandell, 2008). PRF size exhibits significant local variations superposing the expected pattern of pRF size increase with increased eccentricity. Gain and exponent estimates appear largely homogenous with local variations without any systematic pattern. Hemodynamic parameter estimates were largely homogenous close to the prior means.

**Figure 8 (2-column).**
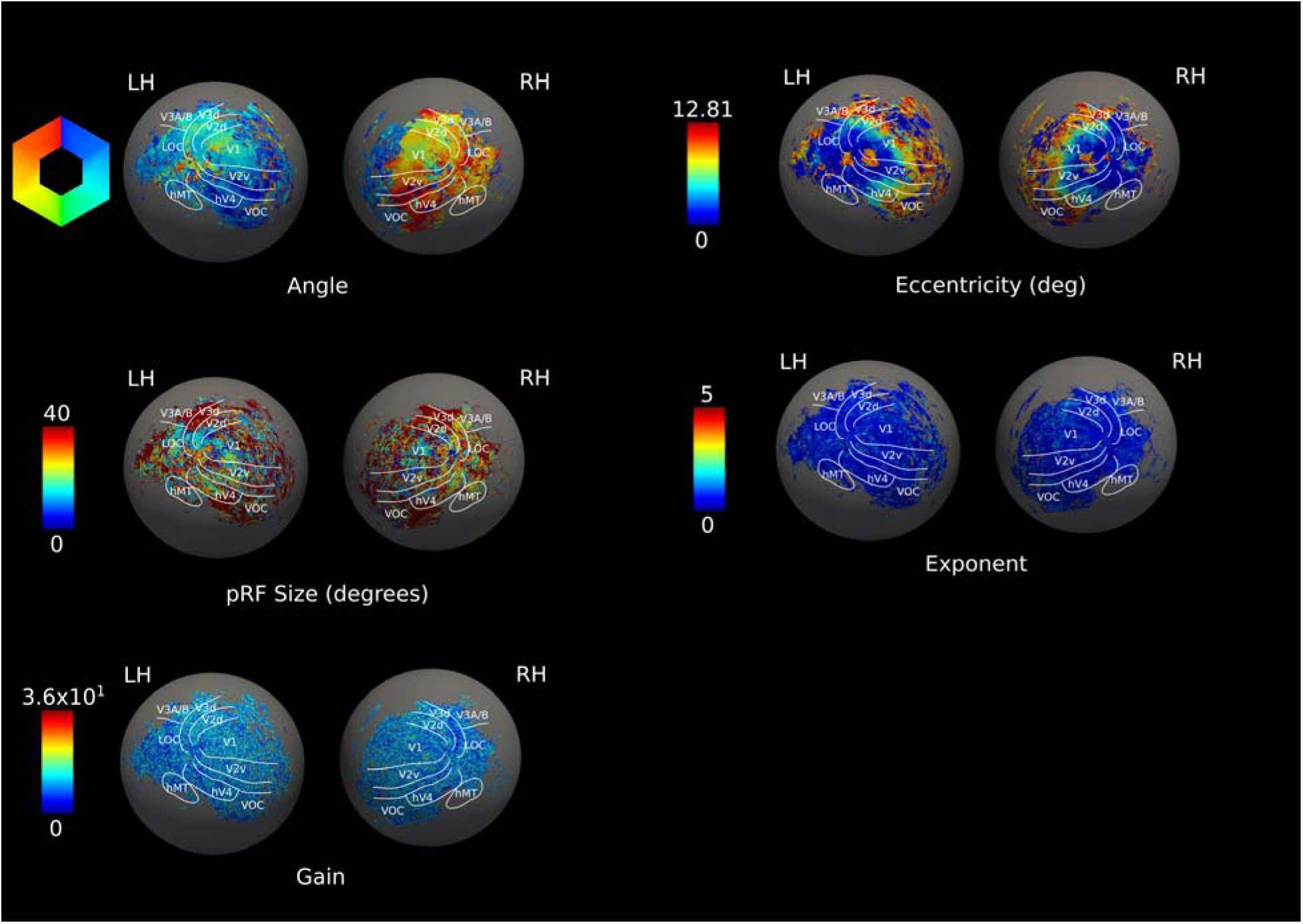
MCMC Parameter estimates (two-column figure). The CSS-pRF parameters Θ, r, σ, expt and gain estimated using MCMC are reported on the inflated left/right hemispheres of a single subject – Subject #1. Two leftmost columns show the parameter means whereas the two rightmost columns show the standard deviations of respective parameters.

### Performance

As opposed to (Adaszewski et al., 2016) where we obtained samples from the posterior distribution for a limited number of points (~1000), in the present study using a new sampling software implementation all ~50000 points in the fMRI dataset were sampled. The runtime of the MH scheme was 48 hours when executed on an NVidia Quadro K2000 GPU (384 computation cores) card which allowed us to perform MH schemes for 384 observation points in parallel. It is a marked improvement over the previously estimated time of ~16 days required to perform MH sampling on all 50000 voxels. The computational efficiency improvements can be fully attributed to the Gaussian premultiplication and interpolation approach. Usage of a lookup table (LUT) eliminates the overhead of computing the exact product of the 2D Gaussian and the visual stimulus mask. This overhead is significant and would dominate the total run time of the GPU code. The main advantage of a LUT is the reduction of dimensionality of data involved in computing the “pRF Gaussian”-related component of the generative model from 3 (stimulus/Gaussian in 2D+time) to 1 dimension (time). In the case of this study - due to usage of the LUT - computing the Gaussian-related component of the model involves just T=734 tri-linear interpolations (where T = duration of the stimulus) for each voxel in each MCMC iteration. Without a LUT this computation would involve 734 evaluations of 2D Gaussian density and multiplications with corresponding stimulus value per each pixel of the stimulus mask. Therefore, with a stimulus size of 100x100 the estimated performance drop without the LUT would be on the order of 10000. It is therefore evident that the LUT brings tremendous performance advantages and could considerably accelerate not only GPU- but also CPU-based samplers. The runtime of RStan’s NUTS sampling scheme for one voxel was 5 hours using the LUT method and 20 days using full Gaussian evaluation, while giving approximately the same results (Table 5).

**Table 5.** Comparison of ground truth, RStan with LUT and RStan without LUT parameter values for one voxel.

|  | **X** | **Y** | **RFSize** | **Expt** | **Gain** | **Kappa** | **Gamma** | **Tau** | **Grubb** | **Rho** |
| --- | --- | --- | --- | --- | --- | --- | --- | --- | --- | --- |
| **Ground Truth** | 19.55 | 13.63 | 35.59 | 0.18 | 23.05 | 0.65 | 0.41 | 0.98 | 0.32 | 0.34 |
| **With LUT** | 19.79 | 13.77 | 23.53 | 0.45 | 21.7 | 0.65 | 0.41 | 1.02 | 0.32 | 0.34 |
| **Without LUT** | 19.25 | 13.77 | 21.01 | 0.38 | 20.08 | 0.66 | 0.41 | 1.01 | 0.32 | 0.34 |

### Tradeoffs of Using a Lookup Table

Regarding the tradeoffs associated with the usage of a LUT heuristic we recognize two types: i. impact on specific behaviors of the MCMC sampler; ii. impact on aggregated metrics such as model log-evidence. General limitations of the method are the following: i. parameter space is bounded by the lookup table; ii. resolution of the lookup table influences accuracy of approximation of the pRF signal; iii. doubling the LUT resolution causes 8-fold increase in lookup table size (frequently seen computation/memory tradeoff – the burden of cubic increase in computational complexity is replaced by cubic increase of LUT size); iv. the assumption about local linearity of the pRF signal form as function of Gaussian parameters is problematic if we want to sample large parameter space because the LUT has to be coarse (due to memory limitations). We believe that the impact of these limitations on specific sampler functions has been observed within this study as abnormalities such as slightly inaccurate estimates of exponent, gain and RF size. Impact of change in both polar angle, radius and RF size parameters is smaller when RF size is big. However it has increasing impact when RF size is small (Figure 4). This is particularly visible when polar angle and radius of the pRF are not perfectly aligned with the ground truth, i.e. with big RF size the signal change is slighter than with small RF size where the peaks will be visibly shifted. Smaller RF size requires the polar angle and radius to match the ground truth much more accurately in order to have good log-evidence and accept a sample. We therefore hypothesize that due to inherent inaccuracy of LUT interpolation RF sizes are positively biased which entails (as compensation) positive bias for gain and exponent as well - since enlarging the Gaussian usually decreases signal amplitude – this is compensated by increased exponent and gain. A few potential solutions come to mind with regards to the above limitations: i. the sampler could be allowed to wander away from parameter bounds covered by the LUT and perform full pRF model computation as long as it stays outside of the LUT; presumably the sampler would spend most of the time within the most probable region (covered by the LUT) therefore this extension would not be significantly detrimental to the performance; ii. quality of interpolation could be improved by using a signal “warping” scheme rather than tri-linear approach; this would imply determining how the signal is “warped” between neighboring cells of the LUT and applying such transformation proportionately to the position of desired parameter point between the LUT cells; to this end we could adapt existing schemes such as Dynamic Time Warping (DWT) (Gupta et al., 1996) or develop new approaches; one potential novelty we considered was usage of a “1.5D” diffeomorphic transform which could be computed from signal change as we follow sub-steps between two LUT cells.

### Tuning the MCMC Sampler

As commonly observed in Monte-Carlo approaches - we believe that recalibration might be necessary depending on the scanning protocol, visual stimulus, as well as chosen hemodynamic (3-/5- parameter Balloon or fixed HRF) and pRF (Dumoulin-Wandell or CSS) components of the model. In the present study we have relied on multivariate effective sample size (mESS) measure (Vats et al., 2015) in order to establish the thinning factor. For the fMRI data, knowing the ESS (ranging from 8 to 50) of a run without thinning we kept increasing the thinning factor as long as the ESS of any thinned series remained within +/- 1 sample. This resulted in the choice of 4 as the thinning factor. We have used the Heidelberg-Welch measure (Heidelberger and Welch, 1983) in a mass-univariate scheme to assess stationarity of the posterior distribution. As burn-in we have arbitrarily chosen to discard half of the samples. We have since reflected upon our thinning approach and our method of choice in the future would be not to use thinning at all (Link et al. 2012; MacEachern et al., 1994; Geyer, 1992) and report ESS measures instead like we do for the simulated data.

### Ambiguities Between pRF and HRF Parameter Estimation

Joint estimation of pRF-HRF parameters is susceptible to ambiguity due to interdependencies between the former and the latter. Bayesian model inversion (either by means of free energy optimization or MCMC sampling) allows to quantify the degree of uncertainty implicated in simultaneous estimation of pRF and HRF parameters by means of the covariance matrix which is absent in point-estimate methods (such as analyzePRF - (Kay et al., 2013)). Given the stochastic nature of MCMC sampling a systematic bias in mean estimates shouldn’t be present unless stemming from or substantiated by further causes. We discuss examples of lookup-table related biases in the “Tradeoffs of Using a Lookup Table” subsection in the Discussion. On the other hand any covariance between pRF and HRF parameters should be reflected in the estimated covariance matrix. Bias in the mean value of gathered samples will be present if a predetermined HRF response is used as the hemodynamic model as opposed to one where HRF parameters are simultaneously sampled and response expressed as a mathematical function. Other sources of estimation uncertainty may include observation noise, subjects’ movement or signal changes not accounted for by used neuronal and hemodynamic models. (Zeidman et al., 2016) include a short discussion of estimation uncertainty and underline that it is important to take into account when making inferences. Using a specifically designed stimulus (e.g. alternating sweep directions - (Harvey and Dumoulin, 2011)) can help to disambiguate between pRF and HRF parameters by reducing the possibility of compensating wrong pRF position with parameters influencing HRF lag. Noteworthy the stimulus used in the present study uses such counterbalanced bar sweeps. The width of pRF remains the most troublesome parameter to disambiguate against HRF width. At this point we believe that this issue remains open and model inversion methods allowing to estimate full covariance matrix allow us to make the most informed inferences by taking into account the estimated covariance between pRF and HRF parameters.

### Future Development: Performance

Current implementation of the MH scheme exhibited satisfactory robustness in testing scenarios. As expected with MCMC approaches, computational load necessary to perform a full sampling run remains the biggest limitation to gathering very large numbers of samples for multiple subjects. Thanks to the advancements presented in this article however, the extension to large datasets becomes more of an inconvenience rather than an impossibility.

To further shorten the execution time, a number of optimizations remain to be tested. Evaluation of hemodynamic state equations is currently performed using full numerical integration and the time step used is that of the scan repetition time (TR) divided by 20; simple improvement in this part of the code involves decreasing the subdivision factor as long as equations remain stable; reduction to 10 would offer 2-fold speed improvement, whereas reduction to 5 would accelerate the process 4-fold. We haven’t thoroughly optimized this parameter and assumed 20 as an arbitrary value.

A more elaborate improvement would be to re-parametrize state equations using Volterra kernels as suggested in (Friston et al., 2003). This approach would allow to eliminate costly integration altogether, however as before a balance between precision and number of Volterra coefficients would have to be carefully evaluated to maintain performance improvement without degrading simulation fidelity.

As the two-step optimization method described by (Dumoulin and Wandell, 2008) with compressive spatial summation extension by (Kay et al., 2013) takes on average 24 hours using CPU implementation we were aiming to stay within the same order of magnitude and with the ensemble of aforementioned techniques we have achieved this goal. We haven’t compared the performance against pure Dumoulin-Wandell classical model.

Furthermore, addressing the shortcomings of the sampler itself could make the algorithm orders of magnitude faster. A comprehensive overview of gradient-free and gradient-based MCMC samplers applied to Dynamic Causal Modelling (DCM) is presented in (Sengupta et al., 2016, 2015). The authors conclude that gradient-based Hamiltonian Monte-Carlo and Langevin Monte-Carlo both offer superior computational performance than Metropolis-Hastings algorithm. Among the gradient-free samplers, single chain adaptive MCMC is highlighted as possessing better efficiency and requiring no tuning. All of the above schemes are good candidates for replacing our current Metropolis-Hastings approach which was chosen primarily due to its simplicity and in order to avoid unexpected interactions with the new heuristic.

Finally, once the validity of chosen pRF model and its relationships with hemodynamic parameters is established using sampling methods, GPU implementation of Variational Bayes approaches such as (Zeidman et al., 2016) should offer much better estimation performance than an MCMC scheme.

### Future Development: Population Receptive Fields

In the study under consideration, visual stimuli comprised wedges, rings and bar patterns similar to (Dumoulin and Wandell, 2008), however the algorithm permits the use of any sequence of images allowing for completely arbitrary stimuli.

A circular pRF model without inhibitory surround was used, however with a Gaussian premultiplication lookup table approach an attempt could be made to model elliptical pRF with inhibitory surround. This would require to perform n-linear interpolation depending on the sophistication of target pRF model and would significantly increase the size of lookup table which might prove to be the limiting factor to this solution. Decomposing different “parts” (e.g. horizontal, vertical, surround, center, etc.) of the pRF Gaussian into separate lookup tables could prove to be sufficient remedy to this instance of the “curse of dimensionality”.

Sampling approaches offer a powerful tool to investigate the ensemble of pRF, hemodynamic as well as other parameters such as anatomical T1-weighted images, Diffusion-Weighted Images (DWI), Voxel Based Quantification (VBQ) results (Draganski et al., 2011), etc. as long as they’re coupled by a common signal generation model. Longitudinal changes as well as function-behavior relationships (Rigoux and Daunizeau, 2015) could also be investigated along the same lines.

### Conclusion

The method presented in this work combines a population receptive field (pRF) model with neuronal hemodynamic state equations and a Markov Chain Monte Carlo (MCMC) sampling approach in order to estimate full Bayesian posterior distributions of visual and hemodynamic parameters based on a sequence of visual stimuli. Thanks to the novel lookup table heuristic we were able to perform large sampling runs on whole-ROI data within a short allotted time. We showed that the proposed heuristic doesn’t introduce significant error to the signal generated in the forward model. We further showed that estimates obtained with and without using the lookup table are consistent. We inspected the log probability with and without the heuristic across many different models and demonstrated that they are consistent. Using in-vivo dataset we verified the correlation of pRF and hemodynamic parameters and concluded that no spatial pattern can be observed which suggests that using a canonical hemodynamic response function for the entire set of observation points as employed by other methods might be justified.

We discussed a number of future directions for the development of our methodology including adaptive equation integration, application of Volterra kernels, elliptical Gaussian field with inhibitory surround using lookup table decomposition, integration of other modalities (anatomical, functional and behavioral) as well as longitudinal component and group studies.

We also proposed that the pRF/hemodynamic sampling approach could be adapted to serve as validation method for Variational Bayes approaches.

Finally, hoping that members of the neuroimaging and vision communities will find our approach useful we provide a stand-alone toolbox for MCMC sampling of pRF/hemodynamic parameter distributions called QPrf. The software is provided along with its source code under the terms of GNU General Public License version 3 (GNU GPLv3) and is available in the following GitHub repository: https://github.com/sadaszewski/qprf.

